# Biophysical and dynamic characterization of a fine-tuned binding of the human Respiratory Syncytial Virus M2-1 core domain to long RNAs

**DOI:** 10.1101/2020.07.22.216952

**Authors:** Icaro P. Caruso, Giovana C. Guimarães, Vitor B. Machado, Marcelo A. Fossey, Dieter Willbold, Fabio C. L. Almeida, Fátima P. Souza

## Abstract

The human Respiratory Syncytial Virus (hRSV) M2-1 protein functions as a processivity and antitermination factor of the viral polymerase complex. Here it is presented the first evidence that hRSV M2-1 core domain (cdM2-1) alone has an unfolding activity for long RNAs, as well as a biophysical and dynamic characterization of the cdM2-1/RNA complex. The main contact region of cdM2-1 with RNA was the α1–α2–α5–α6 helix bundle, which suffered local conformational changes and promoted the RNA unfolding activity. This activity may be triggered by base-pairing recognition. RNA molecules wrap around the whole cdM2-1, protruding their terminals over the domain. The α2–α3 and α3–α4 loops of cdM2-1 were marked by an increase in picosecond internal motions upon RNA binding even though they are not directly involved in the interaction. The results revealed that the cdM2-1/RNA complex originates from a fine-tuned binding, contributing to unraveling interaction aspects necessary to M2-1 activity.

**IMPORTANCE:** The main outcome is the molecular description of a fine-tuned binding of the cdM2-1/RNA complex and the evidence that the domain alone has an unfolding activity for long RNAs. This binding mode is essential in the understanding of the function in the full-length protein. Orthopneumovirus, as the human Respiratory Syncytial Virus (hRSV), stands out for the unique role of M2-1 as a transcriptional antitermination factor able to increase the RNA polymerase processivity.

## INTRODUCTION

Human Respiratory Syncytial Virus (hRSV) is the major causative agent of lower tract respiratory diseases such as pneumonia and bronchiolitis in children worldwide. This virus is also responsible for respiratory morbidity in the elderly and people with compromised immunity and cardiorespiratory disease. The main hRSV-related risk group is children with congenital immunodeficiency, bronchopulmonary dysplasia, heart disease, hypertension, prematurity, and low birth weight (1–3).

Of the 11 proteins encoded by hRSV, two are expressed by the M2 gene, so called M2-1 and M2-2 (1). The proteins of the M2 gene are involved in the assembly of the active form of the ribonucleoprotein (RNP) complex, with the M2-1 acting in transcription as a processivity and antitermination factor, and M2-2 working as a molecular “switch” between transcription and RNA replication (4).

M2-1 prevents the RNA-dependent RNA polymerase (RdRp) complex from dissociating when it reaches inter and intragenic stop codon and, consequently, increasing the transcription efficiency of genes near the 5’ end (5, 6). Blondot and collaborators (2012) hypothesized three possible explanations for acting as a factor that prevents the premature termination of the transcription, thus important for mRNA transcription: i) The M2-1 protein could bind to the nascent mRNA transcript to favor the transcription elongation, either preventing it to re-hybridize to the template or from forming secondary structures, which would destabilize the transcription complex. ii) The polymerase processivity enhancing effect of M2-1 could be due to an increase in the affinity of the polymerase for the genomic RNA template in a sequence non-specific manner. iii) The M2-1 could recognize gene end sequences either on the nascent mRNA or on the RNA template, preventing the release of the polymerase complex from its template and favoring transcription re-initiation at the downstream gene start sequences (7).

The crystal structure of the hRSV M2-1 protein determined by Tanner and collaborators (2014) shows a tetrameric arrangement with the monomers presenting three structurally distinct regions: the zinc finger domain which is composed of N-terminal residues bind to a zinc atom and aids in the transcription process; the oligomerization helix that is responsible for tetramer formation; and the core domain that acts directly on the interaction with phosphoprotein P and RNA (8). The core domain of M2-1 (cdM2-1) is folded into six α-helices, which are structurally arranged in a α1–α2–α5–α6 helix bundle and α3–α4 hairpin (7, 8). Blondot et al. (2012) determined by NMR studies that α2, α5, and α6-helix of the cdM2-1 play a key role in the binding to nucleic acids, and also demonstrate the preference of the core domain for purine-rich RNAs, especially adenine (7).

The crystal structure of the human Metapneumovirus (hMPV) M2-1 protein, which is homologous to hRSV M2-1, presented a distinct conformation for one of the tetramer core domains, which was termed as open conformation (9). In this conformation, one of the tetramer core domains is located far from the rest of the protein due to a rotation in the flexible linker (residues with electronic density absent in the crystal structures) between the oligomerization and core domain. From molecular dynamics (MD) simulations and small angle X-ray scattering (SAXS) experiments, it was characterized a dynamic open-closed conformation equilibrium for hMPV M2-1, and that the closed conformation is structurally similar to the hRSV M2-1 protein. It was also reported that M2-1 closed conformation is prevalent in the presence of RNA, which is stabilized by the simultaneous binding of the nucleic acid molecule to the zinc finger region and to the core domain of the protein (9).

Recently, Molina and coauthors (2018) observed that RNA of 20 nucleotides (20mer RNA) binds with positive cooperativity to two core domains in the hRSV M2-1 tetramer, allowing RNAs to bridge two adjacent RNA binding sites. No cooperativity was reported for the binding of the core domain to 20mer RNA and of M2-1 tetramer to 10mer RNA (10). They showed by circular dichroism (CD) that 20mer RNAs are unfolded upon the formation of its complex with M2-1 tetramer but, in the presence of only core domain, significant structural changes are not detectable. CD approach indicated that the secondary structures content of M2-1 shows no major changes upon the formation of the complex (10). This finding corroborates with the first of the above-mentioned hypotheses proposed by Blondot and colleagues (2012) for the mechanisms by which M2-1 prevents the polymerase from terminating transcription (7). Molina and collaborators (2018) also reported by fluorescence experiments that the association process of nucleic acids with M2-1 tetramer is composed of two steps: the first being a fast phase related to conformational changes of the RNA molecules, and the second being a slow phase related to subtle rearrangements of the M2-1/RNA complex that occur through an induced-fit mechanism. Based on induced-fit mechanism, subtle conformational rearrangements in the complex take place in the protein moiety. Therefore, in the case of M2-1, as CD experiments indicate that no major secondary structure changes occur upon binding, probably loop or hinge motions may reposition the RNA binding domains (core domains), similarly as shown for the hMPV M2-1 protein, where transition from open to closed conformation is observed for cdM2-1 upon RNA association (9, 10).

Taken together, the aforementioned studies characterize the M2-1 tetramer as a flexible platform for RNA recognition, being the core domain the major binding site responsible for the first molecular contact with the nucleic acid molecules. Despite the well-described biological activity (11, 12) and structures of M2-1 (8, 9), and recent clarifications on the possible biochemical mechanism accounting for its antitermination activity (10), molecular details about biophysical-chemical and dynamic features of the interaction between the core domain of M2-1 and RNA still need to be introduced. In this report, an integrated approach with experimental and computational techniques was employed to obtain information at the molecular level on the binding of hRSV cdM2-1 to long RNAs (~80 bases). The fluorescence spectroscopy experiments at different conditions of temperatures, ionic strengths, and pH revealed a key role of the electrostatic interactions in the formation of the complex and that the van der Waals interactions and hydrogen bonds in the bimolecular stabilization, while differential scanning calorimetry (DSC) measurements indicated a slight increase in thermal stability of cdM2-1 upon RNA binding. Nuclear magnetic resonance (NMR) highlighted structural features of the cdM2-1 and its complex with RNA. The chemical shift perturbation (CSP) analysis identified the α1–α2–α5–α6 helix bundle in the core domain as the main interaction region with RNA and, along with thermal susceptibility measurements of the amide hydrogen (^1^H_N_) chemical shifts, pointed out that this bundle helix may undergo local structural changes that extend to the α3–α4 hairpin. The solvent paramagnetic relaxation enhancement (sPRE) measurements showed that cdM2-1 is permanently protected from solvent exposure when complexed with the nucleic acid, while intensity changes in imino proton resonances suggest a protein-induced unfolding of partial formations of RNA base-pairing. ^15^N relaxation data revealed that cdM2-1 dynamics undergo significant changes after interaction with RNA, presenting a decrease in the overall tumbling motion and locally an increase in the flexibility of the α2–α3 and α3–α4 loops of the protein. The structural models of the cdM2-1/RNA complex generated from docking calculations and evaluated by molecular dynamics (MD) simulations corroborated with experimental results and highlighted Lys150 and Arg151 as pivotal residues for the stabilization of the complex. The results presented herein contribute to understand the molecular behavior of the core domain of hRSV M2-1 in solution and upon binding to its biological partner, the RNA.

## RESULTS

### cdM2-1/RNA interaction investigated by fluorescence spectroscopy

The binding of the core domain of hRSV M2-1 to RNA was investigated using fluorescence quenching experiments at different temperatures (15, 25, and 35 °C), salt concentrations (0, 150, and 350 mM of NaCl), and pHs (5, 6.5, and 8). Fig. 1A shows that the fluorescence intensity of cdM2-1 is quenched with increments of RNA concentration, suggesting that the microenvironment of the fluorophores of the protein is affected by the presence of the nucleic acid. The changes of the fluorescence quenching signal (*F*_0_ – *F*)/(*F*_0_ – *F_S_*) as a function of RNA concentration were analyzed using the Eq. (1) at the temperature of 15, 25, and 35 °C (Fig. 1B). The binding isotherm curves provided the values of dissociation constant (*K_d_*) and stoichiometry coefficient (*n_SC_*) for the cdM2-1/RNA complex, which are shown in Table 1. The values of *K_d_* determined for the cdM2-1/RNA interaction are in the nano to micromolar range and the binding affinity revealed to decrease with increasing temperature from 15 to 35 °C. The average value of *n_SC_* (~2.25) at the three investigated temperatures indicates that at least two cdM2-1 binds to one RNA molecule (Table 1).

**Figure 1.**
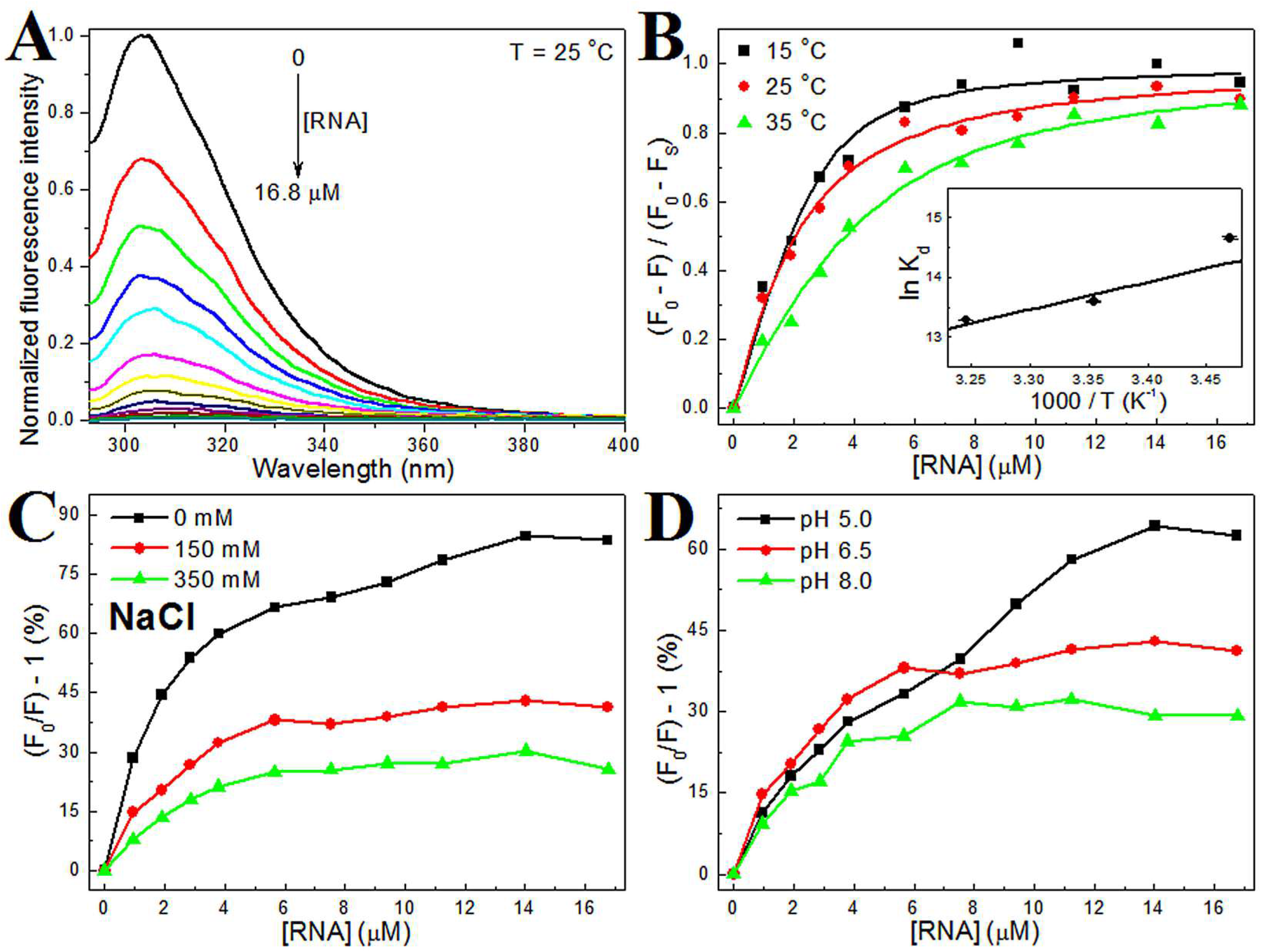
Analysis of the fluorescence quenching data of the cdM2-1/RNA interaction at different conditions of temperature, ionic strength, and pH. (A) Emission spectra of the cdM2-1 in absence and presence of RNA concentration increments (pH 6.5, T = 25 °C, λex = 288 nm). [cdM2-1] = 5.5 μM; [RNA] = 0–16.8 μM. The fluorescence quenching spectra at 15 and 35 °C showed a similar profile to 25 °C (data not shown). (B) The analysis of the fluorescence data of the cdM2-1/RNA interaction. Changes of the fluorescence quenching signal of cdM2-1 as a function of RNA concentrations in 50 mM phosphate buffer (pH 6.5) containing 150 mM NaCl and 1.0 mM DTT at 15, 25, and 35 °C. The insert denotes the van’t Hoff plot used to determine the enthalpy change value in the formation of the cdM2-1/RNA complex. (C, D) Fluorescence quenching percentage rates as a function of RNA concentrations at different conditions of salt concentration (0, 150, and 350 mM NaCl) and pH (5, 6.5, and 8) for the temperature of 25 °C.

**Table 1.**
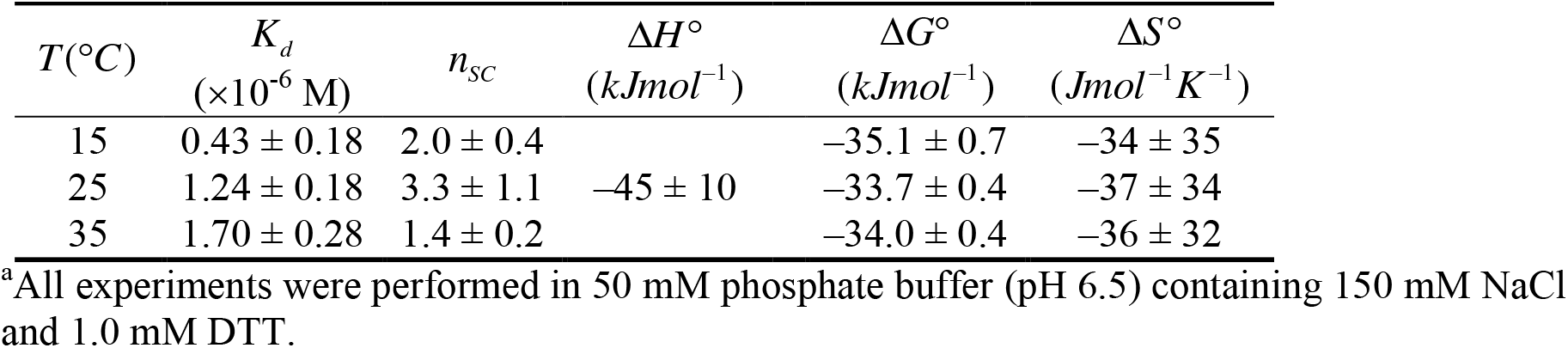
Dissociation constant (*K_d_*), stoichiometry coefficient (*n_SC_*), enthalpy change (Δ*H*), Gibbs free energy change (Δ*G*), and entropy change (Δ*S*) of the cdM2-1/RNA interaction determined using fluorescence quenching experiments at 15, 25, and 35 °C.^a^

| $T(^{\circ}C)$ | $K_d$<br>($\times 10^{-6}$ M) | $n_{sc}$ | $\Delta H^{\circ}$<br>($kJmol^{-1}$) | $\Delta G^{\circ}$<br>($kJmol^{-1}$) | $\Delta S^{\circ}$<br>($Jmol^{-1}K^{-1}$) |
| --- | --- | --- | --- | --- | --- |
| 15 | $0.43 \pm 0.18$ | $2.0 \pm 0.4$ | | $-35.1 \pm 0.7$ | $-34 \pm 35$ |
| 25 | $1.24 \pm 0.18$ | $3.3 \pm 1.1$ | $-45 \pm 10$ | $-33.7 \pm 0.4$ | $-37 \pm 34$ |
| 35 | $1.70 \pm 0.28$ | $1.4 \pm 0.2$ | | $-34.0 \pm 0.4$ | $-36 \pm 32$ |

<sup>a</sup>All experiments were performed in 50 mM phosphate buffer (pH 6.5) containing 150 mM NaCl and 1.0 mM DTT.

The *K_d_* as a function of temperature was analyzed using the van’t Hoff equation [Eq. (2)] (insert in Fig. 1B), which was linear over the investigated temperature range. The enthalpy change (Δ*H*) was −45 ± 10 kJ·mol^−1^, therefore denoting an enthalpically favorable exothermic binding reaction. The values of Δ*G*<0 and Δ*S*<0 indicate that the interaction is spontaneous and entropically unfavorable. The enthalpic term provides the major contribution to the Gibbs free energy change, suggesting that the binding process is enthalpically driven. The negative values of Δ*H* and Δ*S* indicate that van der Waals interactions and hydrogen bonds are important non-covalent interactions responsible for the stabilization of the cdM2-1/RNA complex (13).

Fig. 1C and 1D show the effect of ionic strength (0, 150, and 350 mM NaCl) and pH (5, 6.5, and 8) at 25 °C. With an increase in the salt concentration, the fluorescence quenching percentage rates [(*F*_0_/*F* – 1)%] decreased in each titration step of RNA with cdM2-1 (Fig. 1B), indicating that the rising in ionic strength caused a reduction in the quenching efficiency with tendency to increase the RNA binding constant (1.13 ± 0.11 and 1.70 ± 0.19 μM for *n_SC_* = 2.25 at 0 and 350 mM NaCl, respectively). The decrease in quenching efficiency is probably related to an electrostatic shielding of the protein and/or nucleic acid. The pH change from 8 to 5 in the buffer solution induced an increase in (*F*_0_/*F* – 1)% rates, suggesting that an increment of the acidity promoted an increase in the quenching efficiency of the cdM2-1 by RNA. This fact may be explained as a strengthening of the electrostatic interactions involved in the formation of the cdM2-1/RNA complex, since the ProtParam tool from the ExPASy web server (14) yielded an increase in the total positive charge of the cdM2-1 due to the protonation of His147 and His168 at pH 5. Despite the increase in (*F*_0_/*F* – 1)% rates, there was a decrease in the cdM2-1/RNA affinity (*K_d_* = 0.90 ± 0.14 and 4.04 ± 0.41 μM for *n_SC_* = 2.25 at pH 8 and 5, respectively) due to reduction in pH, which reinforces the interpretation that electrostatic interactions are important for the formation of the complex while hydrogen bonds and van der Waals interactions play a key role in the binding stabilization. Circular dichroism (CD) experiments were used to check the spectral profile of the secondary structures of cdM2-1 at different pHs (Fig. S1). The far UV-CD spectra of the protein at pH 5, 6.5, and 8 presented similar spectral characteristics of α-helix with the minimum of mean residues ellipticity at 208 and 222 nm, indicating that a rise in the alkalinity did not significantly change the profile of secondary structures of the core domain of hRSV M2-1. Therefore, the variations in the fluorescence quenching percentage rates at different conditions of pH did not arise from secondary structure changes.

### Thermal stability of cdM2-1 and cdM2-1/RNA complex determined via DSC

The analysis of the thermal denaturation by differential scanning calorimetry (DSC) obtained over the temperature range of 10–90 °C showed that cdM2-1 has an endothermic transition with a melting temperature (*T_m_*) at 68.7 ± 0.4 °C and a calorimetry enthalpy change (Δ*H_cal_*) of 174 ± 18 kJ.mol^−1^ in 50 mM phosphate buffer (pH 6.5) containing 150 mM NaCl and 1.0 mM DTT (Fig. 2). Interestingly, Teixeira and collaborators (2017) showed a *T_m_* value of ~55 °C for the full-length hRSV M2-1, which is at least 13 °C lower compared to that of its core domain (15). The van’t Hoff enthalpy change (Δ*H_vH_*) calculated from integration process of the cdM2-1 thermogram revealed a value of 341 ± 30 kJ.mol^−1^ that is different from Δ*H_cal_*. This result indicated that the temperature-induced unfolding process of the core domain of hRSV M2-1 did not follow a two-state model. The reversibility test (cooling scan) showed that the thermal denaturation of the protein is irreversible, showing no transition in the cooling process. *T_m_* and Δ*H_cal_* values for the thermal unfolding process of the cdM2-1/RNA complex were 69.4 ± 0.1 °C and 136 ± 10 kJ.mol^−1^ for a protein/RNA molar ratio of 1:0.5, and 69.8 ± 0.1 °C and 103 ± 8 kJ.mol^−1^ for a molar ratio of 1:1, respectively (Fig. 2), indicating that the binding of cdM2-1 to RNA promoted a slight change in thermal stability of the protein.

**Figure 2.**
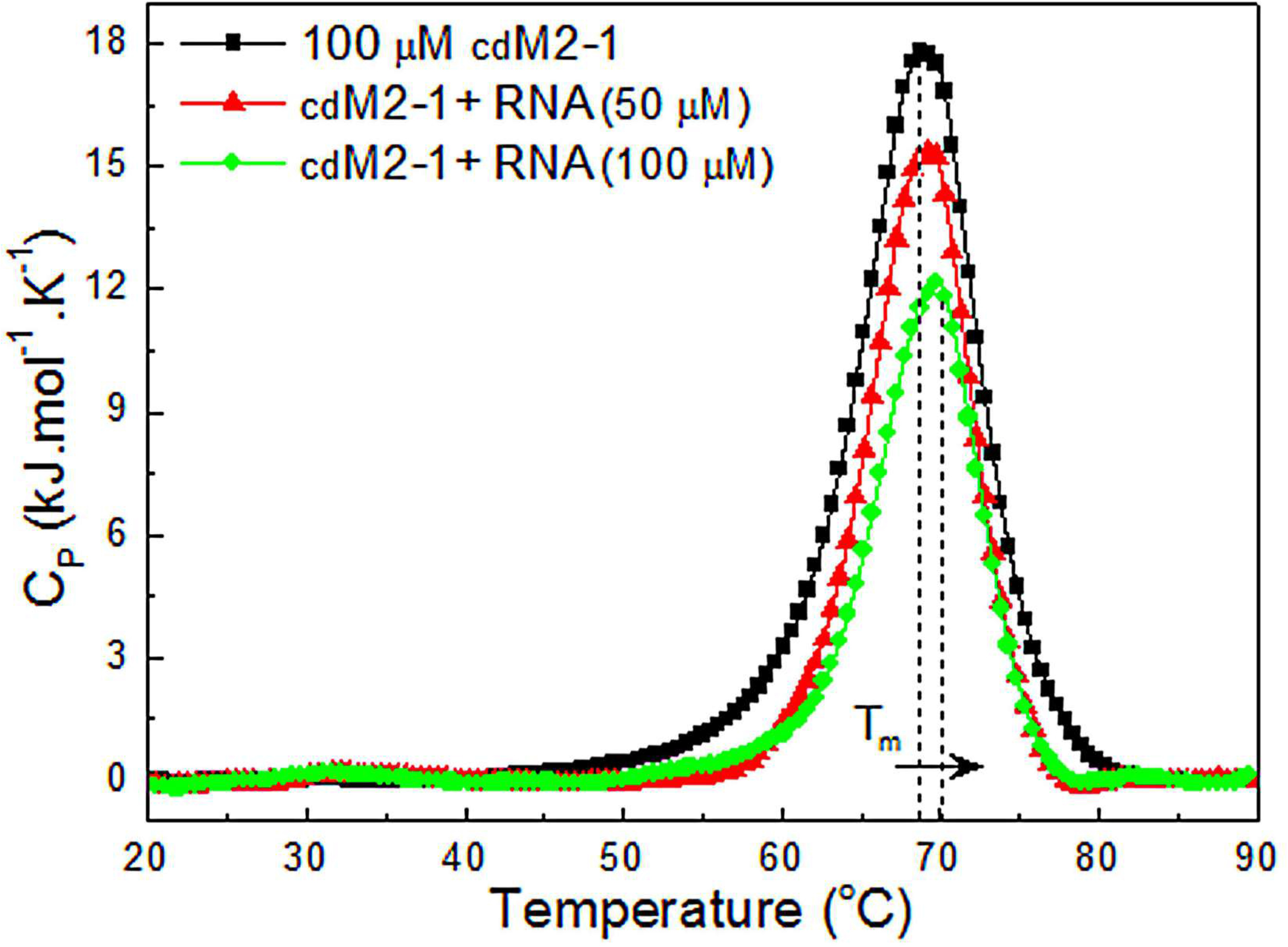
Thermal stability of the free and RNA-bound cdM2-1. DSC thermograms of the protein and its complex with RNA collected in 50 mM phosphate buffer (pH 6.5) containing 150 mM NaCl and 1.0 mM DTT using scan rate of 1.0 °C/min. The dotted lines denote the melting temperature (*T*_m_) for cdM2-1 in absence and presence of RNA (100 μM). The concentration of the protein was of 100 μM and nucleic acid of 0 (black square), 50 (red triangle), and 100 μM (green circle).

### Mapping cdM2-1 interaction with RNA

The analysis of chemical shift perturbation (CSP) by NMR spectroscopy was also used to characterize the cdM2-1/RNA binding. The NMR titrations of [U-^15^N] cdM2-1 with nucleic acid were carried out using the assigned 2D ^1^H–^15^N HSQC spectra of the free and RNA-bound protein (Fig. 3A). The CSP experiments presented a fast chemical exchange process between free and bound forms on the NMR chemical shift timescale (insert in Fig. 3A). The amino acid residues associated with resonances that showed chemical shift perturbation Δ*δ*(^1^H,^15^N) higher than the average value plus standard deviation between free and RNA-bound cdM2-1 at 25 °C were [arrows in Fig. 3A, Δ*δ*(^1^H,^15^N) > average (Δ*δ_ave_*) plus standard deviation (SD) in Fig. 3B, green residues in Fig. 4A]: N-terminal (Leu74), α2-helix (Lys92, Gln93 side chain, Val97), α5-helix (His147, Lys150, Arg151), α5–α6 loop (Leu152), and α6-helix (Asp155, Val156). These residues of the cdM2-1 promoted the binding to RNA either by direct interaction or by conformational rearrangements remote from the interaction interface. Fig. S2 presents the arrangement of the residues with significant Δ*δ* values on the full-length hRSV M2-1 structure in the close and open conformation. The residues Gly62, Ala63, Lys92, Gln93, Leu152, and Ala154 presented an increase in line width with increment of the RNA concentration and consequently a decrease in peak intensity and/or disappearance of the resonance under binding to nucleic acid. The chemical shift changes of the amide group of [U-^15^N] cdM2-1 under binding to RNA revealed that the resonances of regions in α1–α2–α5–α6 helix bundle presented the most significant Δ*δ*(^1^H,^15^N) values (cyan and green residues in Fig. 4A). These regions denote a positive electrostatic potential surface (Fig. 4B) which is favorable for the interaction with a negative charge of the phosphate groups from nucleic acid.

**Figure 3.**
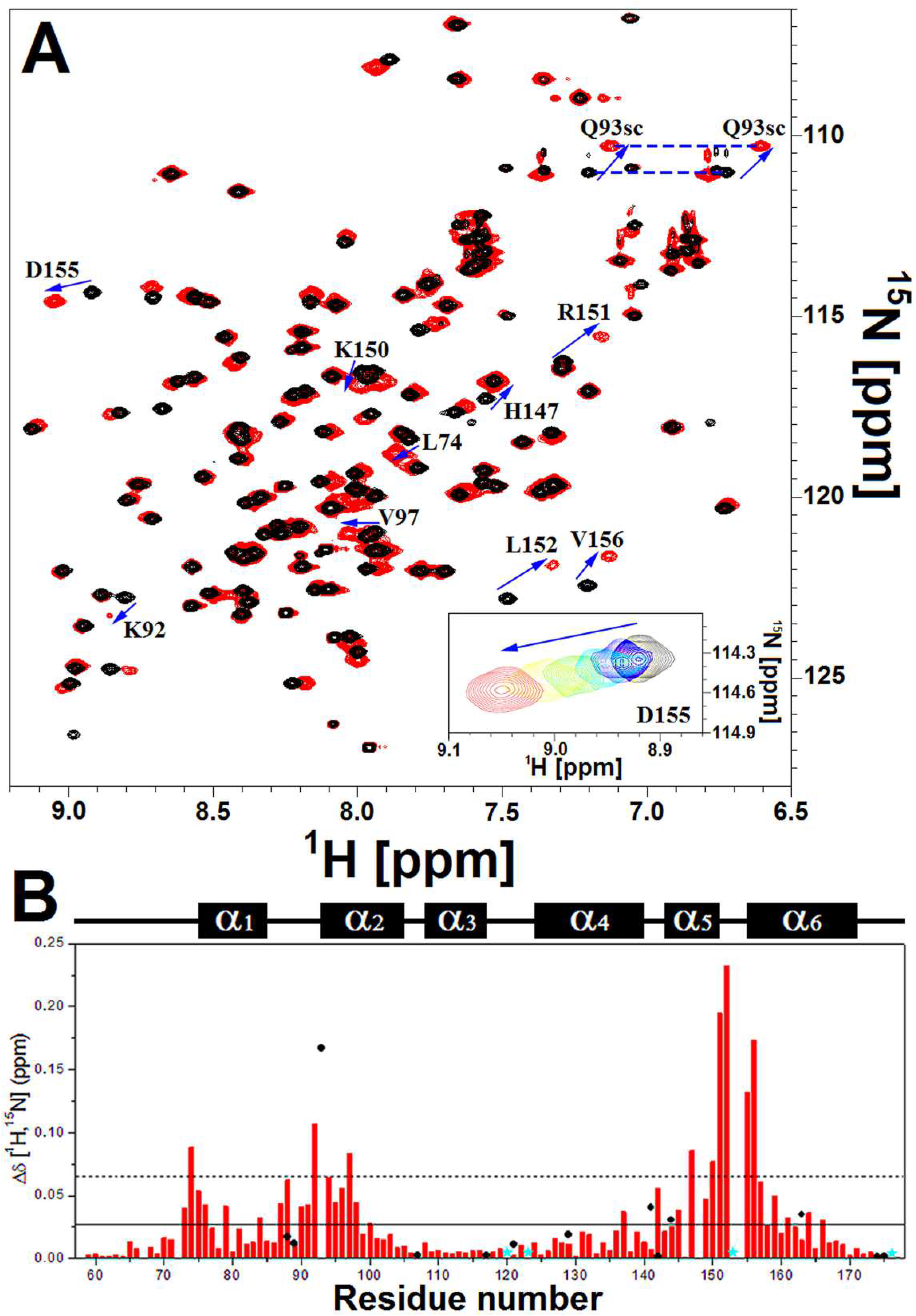
cdM2-1/RNA binding investigated by NMR spectroscopy. (A) Two-dimensional ^1^H–^15^N HSQC spectra of the free (black) and RNA-bound [U-^15^N]cdM2-1 (red) collected by using NMR spectrometer of 14.1 T (^1^H frequency of 600 MHz) at temperature of 25 °C. The arrows indicate the residues that presented a chemical shift perturbation upon RNA binding higher than Δ*δ_avg_* + SD. The top insert presents the titration effect in the behavior of fast exchange regime for the Asp155 upon RNA binding. The spectra were recorded at protein concentration of 100 μM (black) and at RNA concentration of 15 (blue), 30 (cyan), 55 (green), 80 (yellow), and 115 (red). (B) Chemical shift perturbation (Δ*δ*) for the formation of the [U-^15^N]dgM2-1/RNA complex, 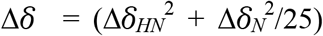. The solid line denotes Δ*δ_avg_* and the dashed line indicates Δ*δ_avg_* + SD. The black circles are Δ*δ* values of side chain of the residues Asn and Gln. The cyan stars indicate the proline residues (Pro120, Pro123, Pro153, and Pro176). The secondary structures along the sequence are indicated at the top.

**Figure 4.**
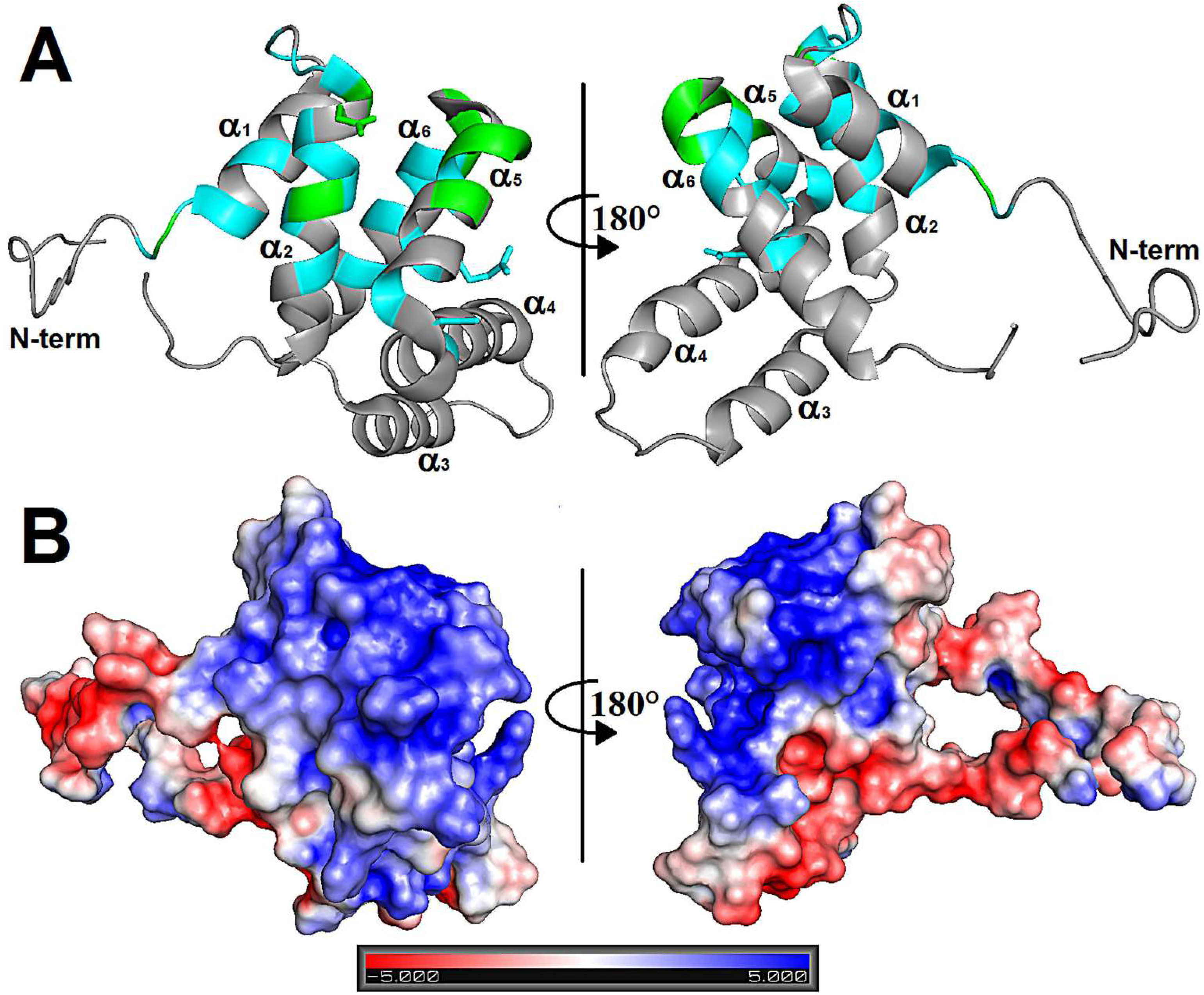
RNA-binding site of cdM2-1 determined by NMR spectroscopy. (A) Identification on the cdM2-1 structure of residues that present significant chemical shift perturbation (Δ*δ*) upon RNA-binding. Values of Δ*δ* higher than Δ*δ_ave_* are colored in cyan: Ala73, Gly75, Val76, Val79, Ile84, Ile87, Asn88, Ile90, Thr91, Ser94, Ala95, Cys96, Ala98, Ser100, Asn142, Thr145, Leu149, Leu157, Lys158, Lys159, Ile161, and Thr164; and higher than Δ*δ_avg_* + SD are colored in green: Leu74, Lys92, Gln93, Val97, His147, Lys150, Arg151, Leu152, Asp155, and Val156. (B) Electrostatic potential surface of cdM2-1 calculated from APBS software using the charge values and protonation states (pH 6.5, 150 mM NaCl, 25 °C) determined by PDB2PQR webserver along with PROPKA program. The bar denotes the electrostatic potential range from −5 (red) to +5 kT (blue).

### Solvent PRE for free and RNA-bound dgM2-1

Solvent exposure measures of each cdM2-1 backbone amide group and its complex with RNA were obtained from solvent paramagnetic relaxation enhancement (sPRE) experiments (Fig. 5). For the protein in the absence of RNA, it is possible to observe that the greatest sPRE effects occur in the N and C-terminus of cdM2-1 and mainly in the N-terminal residues of α1-helix, characterizing these regions as significantly exposed to the solvent. Significant sPRE effects are also observed in virtually all C-terminal residues of the protein loop regions, highlighting the loop between α1 and α2, named α1–α2 loop (Fig. 5). In general, the helical secondary structure regions have the lowest sPRE effects, indicating that the amide groups of the residues in these regions are protected from solvent exposure. An exception to this is α4-helix which exhibited an *i* + 4 pattern of solvent-exposed residues (Ser122, Arg126, Thr130, Ile133, and Ser137) corresponding to a nearly α-helix turn (Fig. S3).

**Figure 5.**
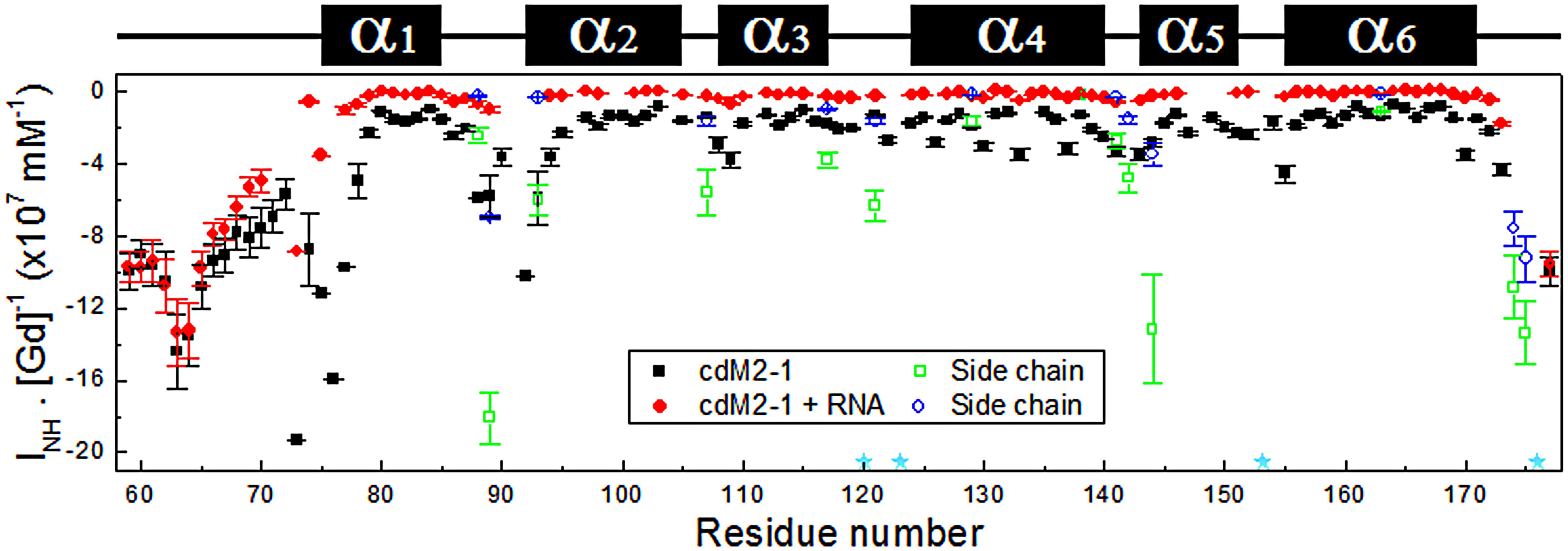
Solvent paramagnetic relaxation enhancement for free and RNA-bound cdM2-1. Solvent exposure (I_NH_·[Gd]^−1^) of each backbone NH amide of the protein (black square) and its complex with RNA (red) measured as a function of the residue number. [cdM2-1] = 350 μM and [RNA] = 115 μM, 25 °C, and 14.1 T (^1^H frequency of 600 MHz). The green squares and blue circles denote the solvent exposure for side chain of the residues Asn and Gln. The cyan stars indicate the proline residues (Pro120, Pro123, Pro153, and Pro176). The secondary structures along the sequence are indicated at the top.

For the cdM2-1/RNA complex, it can be seen a significant reduction of the sPRE effect throughout all the residues of the protein, except the N and C-terminus (Fig. 5). This result indicates that the helical secondary structure regions, along with their loops, are permanently protected from solvent exposure when cdM2-1 is complexed with nucleic acid, while the terminals of the protein remain exposed to the solvent as observed for free cdM2-1. Thus, N and C-terminus of the core domain of hRSV M2-1 might be considered as intrinsically negative control regions for the performed experiments.

### Thermal susceptibility measurements of the amide hydrogen chemical shifts

The thermal susceptibility measurements of the amide hydrogen (^1^H_N_) chemical shifts (*dδ_HN_/dT*) of cdM2-1 in absence and presence of RNA were obtained from 2D ^1^H–^15^H HSQC spectra at 15, 20, 25, 30, and 35 °C (Fig. 6A, see Materials and Methods for details about the interpretation of *dδ_HN_/dT* values). In general, cdM2-1 residues taking place in secondary structure elements presented *dδ_HN_/dT* values higher than −5.0 ppb·K^−1^, excepting Ile146 (green sticks in Fig. 6C) in α5-helix. On the other hand, most values of *dδ_HN_/dT* lower than −5.0 ppb·K^−1^ were reported for residues in the N and C-terminus and loops between helices of the core domain. However, Glu59, Ala63, Asp67, Thr69, Glu70, Glu71, and Ala73 in N-terminal of the cdM2-1 showed *dδ_HN_/dT* > −5.0 ppb·K^−1^, indicating that the amide group of these residues might form less expandable hydrogen bonds (correlated to stronger hydrogen bonds), typically as it is observed for ^1^HNs in secondary structures (16).

**Figure 6.**
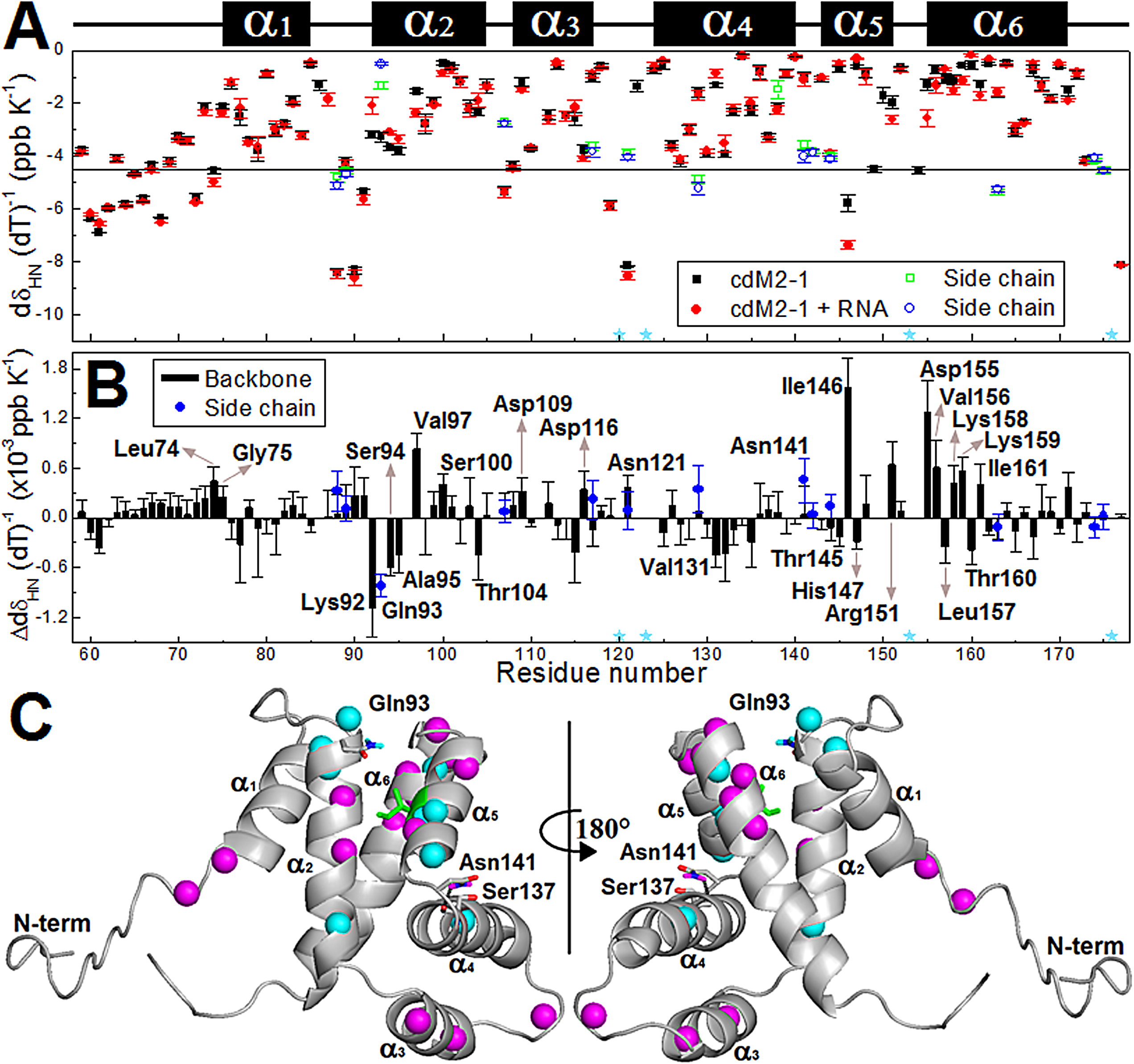
Thermal susceptibility data of the amide hydrogen (^1^HN) chemical shifts of the free and RNA-bound cdM2-1. (A) *dδ_HN_/dT* values of the protein (black square) and its complex with RNA (red circle) as a function of the residue number. The temperature dependence measurements were obtained from 2D ^1^H–^15^H HSQC spectra at 15, 20, 25, 30, and 35 °C. [cdM2-1] = 350 μM and [RNA] = 115 μM, and 14.1 T (^1^H frequency of 600 MHz). The green squares and blue circles denote the values of *dδ_HN_/dT* for side chain of the residues Asn and Gln. The cyan stars indicate the proline residues (Pro120, Pro123, Pro153, and Pro176). The secondary structures along the sequence are indicated at the top. (B) Difference between the values of *dδ_HN_/dT* for the free and RNA-bound cdM2-1 as a function of the residue number 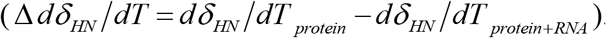. Δ*dδ_HN_/dT* + SD values far away from the zero are considered as significant changes, and the correspondent residue names are denoted. (C) The significant changes of |*dδ_HN_/dT*| values between the free and bound states of the cdM2-1 are indicated in the protein structure. The values of Δ*dδ_HN_/dT* >0 (increase in |*dδ_HN_/dT*|) and Δ*d δ_HN_/dT* <0 (decrease in |*dδ_HN_/dT*|) of the ^1^HN chemical shifts are denoted as cyan and magenta spheres, respectively. The side chain of Gln93 and Asn141 are displayed as sticks with amide hydrogens colored according the change of Δ *dδ_HN_/dT*. The side chain of Asn141 forms a hydrogen bond (black dashed line) with backbone of Ser137. Ile146 is shown as green sticks because presents *dδ_HN_/dT* > −5.0 ppb·K^−1^.

The binding of the cdM2-1 to nucleic acid promoted significant changes of *dδ_HN_/dT* values for residues in N-terminal (Leu74), α1 (Gly75), β2 (Lys92, Gln93 side-chain, Ser94, Ala95, Val97, Ser100, Thr104), α3 (Asp109, Asp116), α3-α4 loop (Asn121), α4 (Val131), α4-α5 loop (Asn141 side chain), α5 (Thr145, Ile146, His147, Arg151), and α6 (Asp155, Val156, Leu157, Lys158, Lys159, Thr160, Ile161) (Fig. 6A and 6B; Δ*dδ_HN_/dT* + SD values far away from the zero). Most of these residues map the protein/RNA interaction interface delimited by regions in the N-terminal, 1, 2, 5, and 6 (Fig. 6C), which corroborates with the results of the CSP experiments (Fig. 3 and 4). Leu74, Gly75, Val97, Ser100, Asp109, Asp116, Asn141 side-chain, Ile146, Arg151, Asp155, Val156, Lys158, Lys159, and Ile161 showed an increase in |*dδ_HN_/dT*| (Δ*dδ_HN_/dT* > 0) which indicates that hydrogen bonds formed by ^1^HN of these residues became more expandable, possibly weaker; while hydrogen bonds stablished by ^1^HN of Lys92, Gln93 side-chain, Ser94, Ala95, Thr104, Val131, Thr145, His147, Leu157, and Thr160 turned into less expandable since the values of |*dδ_HN_/dT*| decreased (Δ*dδ_HN_/dT* <0). In the cdM2-1/RNA binding interface, there are two clusters of residues with their ^1^HN forming hydrogen bonds that became less and more expandable, being the first cluster composed of residues Lys92, Gln93 side-chain, Ser94, and Ala95 in the N-terminal of α2-helix and the second one, formed by Asp155, Val156, Lys158, Lys159, and Ile161 in the N-terminal of α6-helix.

### Evidence of cdM2-1-induced structural changes in RNA

The one-dimensional (1D) ^1^H NMR spectra of protein/nucleic acid complexes reveal that most of the resonance signals from the nucleic acid and protein are overlapped, as it can be observed from amine/amide proton region (6–10 ppm) in Fig. 7. The only spectral region that is not disturbed by the protein resonances is the imino proton region from 11 to 15 ppm (17). The resonances of imino protons are directly involved in base-pairing, which is remarkable evidence of the secondary structure organization of the nucleic acid. Fig. 7 shows the imino proton signals of the protein-free and bound forms of the RNA, as well as the spectrum of the free cdM2-1 as a negative control since proteins have no observable resonances at this spectral region. The 1D ^1^H spectrum of the protein-free RNA presented resonance signals of imino protons at 11–15 ppm range (blue line, Fig. 7), which indicates a structural arrangement level due to partial formations of RNA base-pairing. On the other hand, the ^1^H spectrum of the cdM2-1-bound RNA showed a significant decrease in the intensities of the imino proton signals (red line, Fig. 7), which suggests structural changes in RNA molecules due to its interaction with the core domain, such as a protein-induced unfolding process. No significant line broadening was observed in the amine/amide region (6–10 ppm, in Fig. 7) of the 1D ^1^H spectra, which shows that the decrease in the intensities of the imino proton signals is not due to the larger size of the cdM2-1/RNA complex.

**Figure 7.**
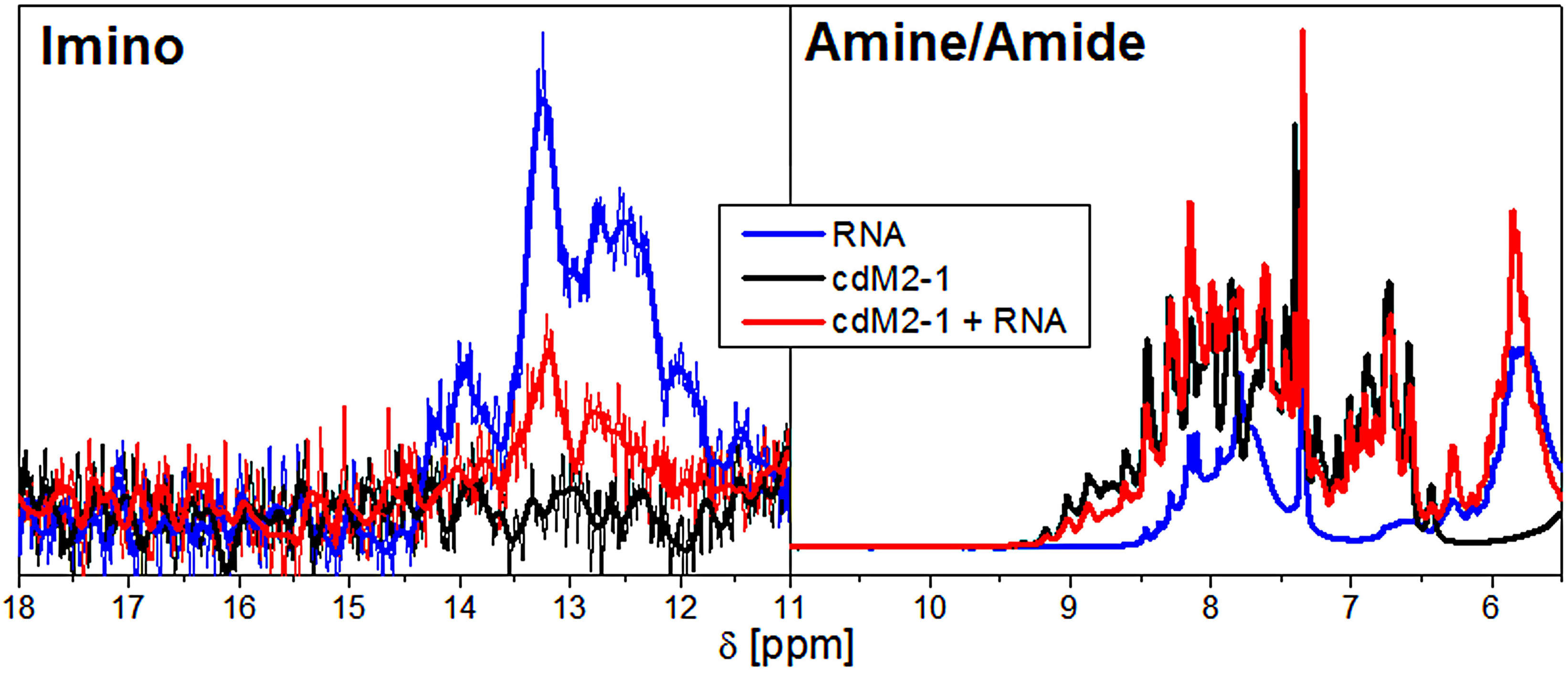
Evidence of the structural changes of RNA induced by cdM2-1. 1D ^1^H NMR spectra of the imino (11–15 ppm) and amine/amide (6–10 ppm) proton region of the protein-free (blue line) and bound (red line) state of RNA. The black line presents the 1D ^1^H spectrum of the free cdM2-1 as a negative control, especially for the imino region. [cdM2-1] = 350 μM and [RNA] = 115 μM, 25 °C, and 14.1 T (^1^H frequency of 600 MHz). In imino proton region, the thick lines denote smoothed profiles that assist the visualization of the spectra.

### Backbone dynamics of the hRSV cdM2-1 and its complex with RNA

The ^15^N relaxation data of the protein backbone by NMR on pico to nanosecond timescales revealed the molecular dynamics of the protein in absence and presence of RNA. The results of the spin nuclear relaxation experiments with the cdM2-1 and cdM2-1/RNA complex were compared and shown in Fig. S4. The ^15^N R_1_, R_2_, and hetNOE relaxation parameters of the core domain presented significant changes in the presence of RNA, except for the N and C-terminal regions of the protein. The lowest values of R_2_ and hetNOE (some of them negative) for N and C-terminal residues of the cdM2-1 characterize these regions as flexible even after the formation of the complex with RNA, thus suggesting terminal residues are not involved in the interaction. The average R_1_, R_2_, hetNOE values excluding the N and C-terminal residues for the free core domain were 1.42 ± 0.05 s^−1^, 11.32 ± 1.10 s^−1^, and 0.76 ± 0.05 units, respectively; whereas the corresponding values for the RNA-bound cdM2-1 were 1.09 ± 0.05 s^−1^, 22.24 ± 2.07 s^−1^, and 0.76 ± 0.11 units, respectively (Table S1).

The reduced spectral density mapping provides insight into internal motions of a protein with no *a priori* assumptions about its diffusion model. The spectral density functions *J*(0.87*ω_H_*), *J*(*ω_N_*), and *J*(0) are sensitive to overall and intramolecular motions on pico to nanosecond timescales. *J*(0.87*ω_H_*) and *J*(*ω_N_*) report fast motions at picoseconds while *J*(0) probes motions at nanoseconds, with possible contributions from the chemical or conformational exchange term. The values of *J*(0.87*ω_H_*), *J*(*ω_N_*), and *J*(0) for the free and RNA-bound cdM2-1 were calculated from Eqs. (3) – (5) and are presented in Fig. 8. As with the relaxation parameters, the spectral density values of the cdM2-1 showed outstanding changes in presence of RNA, except for the N and C-terminal of the protein. The highest and lowest values of *J*(0.87*ω_H_*) and *J*(0), respectively, for residues in the N and C-terminal of the protein indicate that these regions are undergoing fast internal motion on the picosecond timescale and are highly flexible even after the binding to the nucleic acid. The spectral density functions in Fig. 8 and correlation plots of *J*(0.87*ω_H_*) & *J*(0) and *J*(*ω_N_*) & *J*(0) in Fig. S5 revealed a different dynamic profile for the N and C-terminals of the free and RNA-bound cdM2-1 from the protein helical-core region. The plots of *J*(0.87*ω_H_*) & *J*(0) (Fig. S5A and S5C) indicated that these two spectral densities were significantly correlated with correlation coefficient of −0.934 and −0.931 for the cdM2-1 in absence and presence of RNA, respectively. On the other hand, *J*(*ω_N_*) & *J*(0) plots (Fig. S5B and S5D) showed lower correlation coefficients than the *J*(0.87*ω_H_*) & *J*(0) pair with values of 0.642 and −0.227 for the RNA-free and bound forms of the protein, respectively. It is noteworthy that before formatting the bimolecular complex, the plot of *J*(*ω_N_*) & *J*(0) presented a positive correlation and after that, a negative correlation. The average values of *J*(0.87*ω_H_*), *J*(*ω_N_*), and *J*(0) excluding the terminal residues for the free core domain were 5.2 ± 1.0 ps, 258 ± 10 ps, and 3.0 ± 0.3 ns, respectively; whereas the corresponding values for the RNA-bound cdM2-1 were 4.1 ± 1.6 ps, 198 ± 10 ps, and 6.1 ± 0.5 ns, respectively (Table S1).

**Figure 8.**
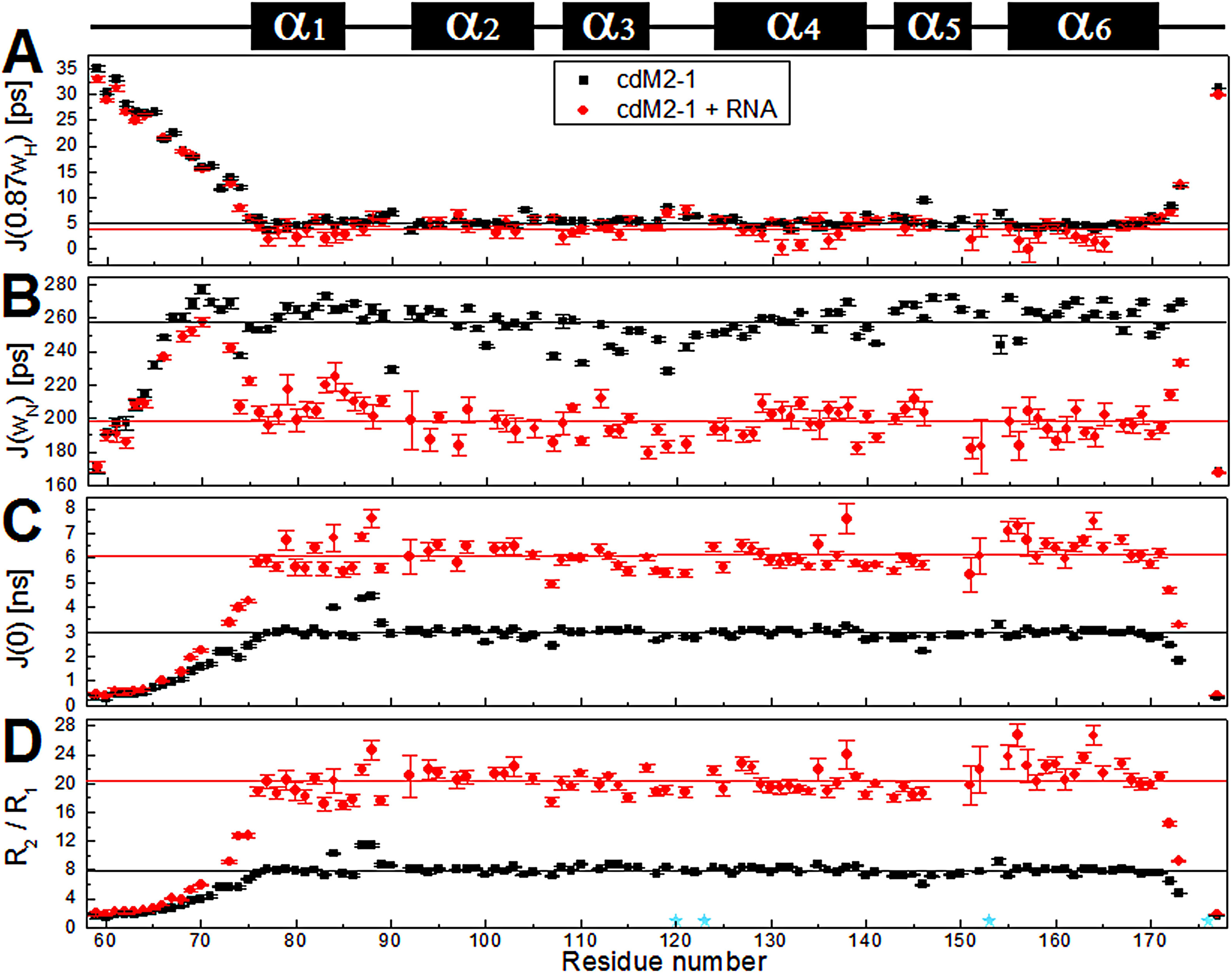
^15^N nuclear spin relaxation data of the free and RNA-bound cdM2-1 backbone amide by NMR on pico to nanosecond timescale. Reduced spectral density function (A) *J*(0.87*ω_H_*), (B) *J*(*ω_N_*), and (C) *J*(0) of the cdM2-1 (black square) and its complex with RNA (red circle) which were calculated from the ^15^N R_1_, R_2_, and hetNOE relaxation data. (D) R_2_/R_1_ ratio calculated from relaxation rates R_1_ and R_2_. [cdM2-1] = 350 μM, [RNA] = 115 μM, 25 °C, and 14.1 T (^1^H frequency of 600 MHz). The black and red lines denote the average values of *J*(0.87*ω_H_*), *J*(*ω_N_*), and *J*(0) for the free and RNA-bound cdM2-1, respectively. To determine the average values, those corresponding to the N and C-terminal residues were excluded. The cyan stars indicate the proline residues (Pro120, Pro123, Pro153, and Pro176). The secondary structures along the sequence are indicated at the top.

The R_2_/R_1_ ratios calculated for all residues of the cdM2-1 in absence and presence of the nucleic acid are shown in Fig. 8D. The average R_2_/R_1_ value excluding the N and C-terminal residues for the free protein was 8.0 ± 0.8 ns, whereas the corresponding value for the RNA-bound cdM2-1 was 20.4 ± 2.2 ns. For the free cdM2-1, the R_2_/R_1_ values obtained for the residues Ile84, Ile87, and Asn88 in α1-helix and α1–α2 loop were above the average, suggesting that these participate in conformational exchange processes (Fig. 8D). After the formation of the cM2-1/RNA complex, the residues Ile87 and Asn88 in α1–α2 loop still presented R_2_/R_1_ values higher than the average while others such as Asp155, Val156, Asn163, and Thr164 in α6-helix had an increase in ratio values above the average.

The extended model-free formalism of Lipari-Szabo was used to analyze the ^15^N R_1_, R_2_, hetNOE nuclear spin relaxation data using TENSOR2 program (18) (Table S2). For the free cdM2-1, the rotational correlation time (*τ_C_*), order parameters (*S*^2^) and exchange rates (*R_ex_*) were obtained from the R_2_/R_1_ ratios. A rotational correlation time for the protein tumbling of 8.53 ± 0.07 ns at 25 °C was calculated using a fully anisotropic model for molecular rotational diffusion (see details in Material and Methods). For comparison with experimental data, the estimation of the *τ_C_* value of the cdM2-1 was calculated using HYDRONMR program (19). It was employed 20 NMR structures from the PDB code access 2L9J (7), excluding amino acid residues from the flexible terminals, and the calculated rotational correlation time was 8.1 ± 0.3 ns at 25 °C. The values of *S*^2^ < 0.7 for the N and C-terminal residues indicated significant thermal fluctuations and high degree of internal mobility for these protein regions (Fig. S6A). On the other hand, some residues located in α1–α2 loop (Ile90), α3-helix (Asp110, Lys113, Leu114), α3–α4 loop (Glu119, Asn121, Ser122), α4–α5 loop (Asn141), and α5-helix (Ile146) showed significantly lower *S*^2^ values than the helical-core average (0.91 ± 0.04, excluding terminal regions), which suggests a certain level of flexibility (Fig. S6C). Conformational exchange contributions with *R_ex_* > 1.0 Hz were observed for the residues Glu59, Ile84, Ile87, Asn88, Ile90, Glu119, and Ala154 of the cdM2-1 (Fig. S6B and S6D). The model-free analysis of Lipari-Szabo for the RNA-bound cdM2-1 did not provide a satisfactory result.

### Computational approach of the cdM2-1/RNA interaction

The structural models of the cdM2-1/RNA complex were generated using the 3dRPC webserver (20). The structural restrains of the complex were defined from the chemical shift perturbations due to the NMR titrations of [U-^15^N]cdM2-1 with RNA. The residues with Δ*δ* > Δ*δ_average_* from NMR CSP analysis were defined as involved in the binding interface. Fig. 9A shows 10 structural models of the cdM2-1/RNA interaction determined by independent docking calculations, which correspond to the lowest energy structures from 10 complex models predicted by the 3dRPC-Score function (20). The 10 RNA molecules crowded around the α1–α2 and α5–α6 loops protruding like an “umbrella” over the α1, α2, α5, and α6 helices and the solvent accessible cleft formed between α1–α2–α5–α6 helix bundle and α3–α4 hairpin. Such behavior can be evidenced by the analysis of the mass density map of the protein-bound RNA (Fig. 9C) and by the change in absolute accessible surface area (Δ*ASA*) of the RNA-bound cdM2-1 (Fig. 9D). The docking calculations also showed that the cdM2-1 preferably binds close to the secondary structure regions of the RNA molecules, where the nucleic acids presented a conformational organization level due to the base-pairing. This fact is exemplified by the cdM2-1/RNA complex model in Fig. 9B, where the α1–α2–α5–α6 helix bundle of the protein lies between two stem-loop secondary structures of the nucleic acid. The evaluation of the non-covalent interactions carried out with PLIP server (21) revealed that the residues Lys150 and Arg151 were involved in salt bridges and hydrogen bonds in nine and eight of 10 cdM2-1/RNA complex models, respectively (Table S3–S12). In general, Lys150 formed two salt bridges with phosphate groups of adjacent nucleotides, while Arg151 established hydrogen bonds with nitrogenous bases and ribose groups of the nucleic acids. These two residues also took place in π-cation interactions with the heterocyclic rings of nitrogenous bases at least three of 10 structural models of the cdM2-1/RNA complex.

**Figure 9.**
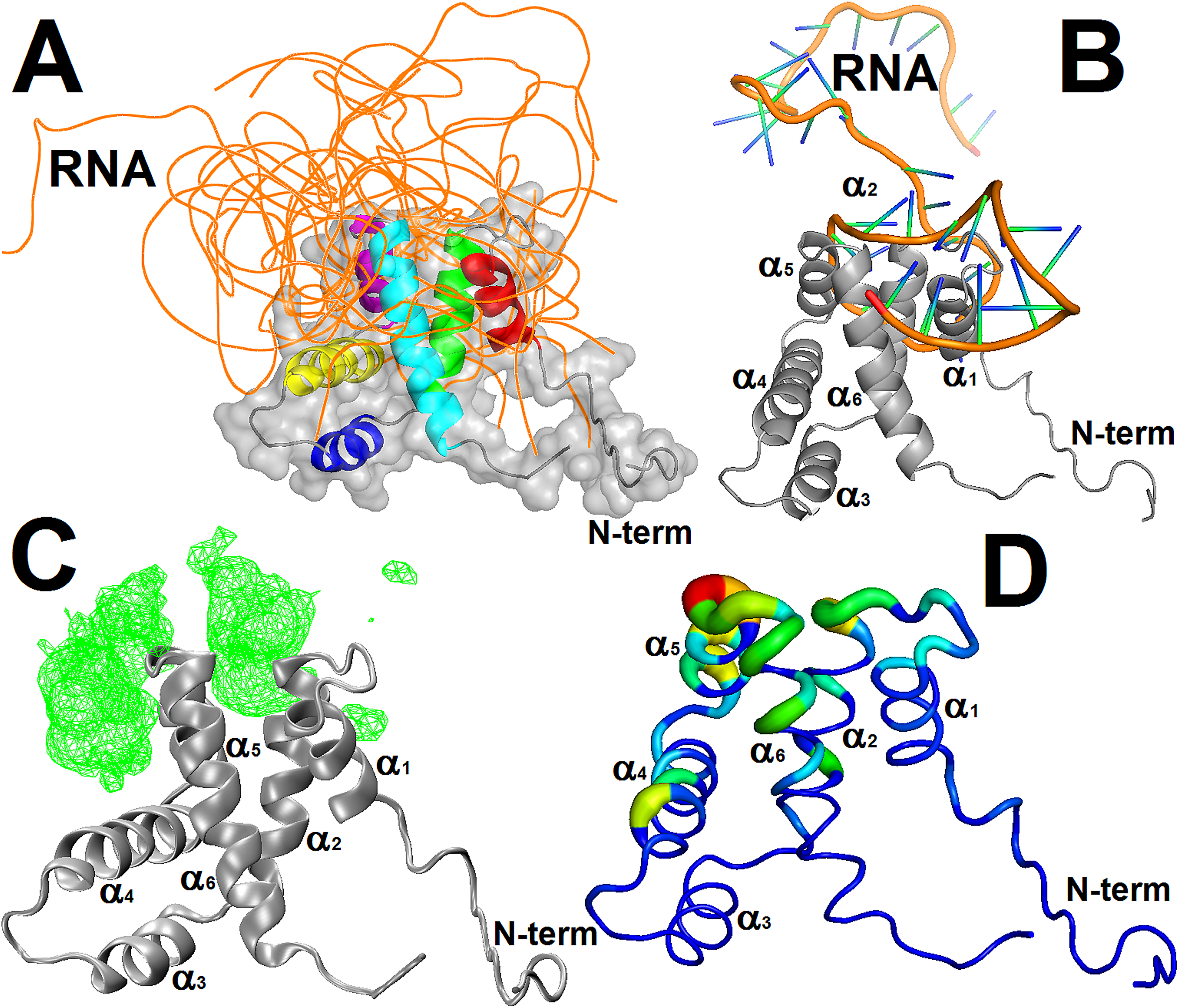
Analysis of the molecular dockings of the structural models of the cdM2-1/RNA complex. (A) Structural models of the cdM2-1/RNA complex calculated by 3dRPC webserver. The backbone of the protein (colorful) and 10 nucleic acids (orange) are presented as cartoon and ribbon, respectively. The protein also is shown as surface representation and its helices are highlighted using different colors (α1: red, α2: green, α3: yellow, α4: blue, α5: purple, α6: cyan). (B) Structural model of cdM2-1/RNA complex which exemplified the preferential binding of the protein (as grey cartoon) close to secondary structural regions of the RNA molecules (as cartoon: backbone in orange and nucleoside in green/blue). (C) Mass density map (green mesh with isovalue of 0.5) of 10 RNA molecules bound to the protein (as grey cartoon) generated by VMD programs using the VolMap tool. (D) Absolute accessible surface area change (Δ*ASA*) of the cdM2-1 (as colorful worm) bound to the nucleic acids. The thickness and colors represent the degree of Δ*ASA*, with the strongly and weakly accessible residues colored in blue (thin) and red (thick), respectively. The structural representations were made in PyMol and VMD programs.

The MD calculations were used to check the structural stability of cdM2-1/RNA complex models generated from the 3dRPC server. The number of hydrogen bonds, number of contacts, and RMSD analyzed from the 20 ns MD simulations for the complex models are presented as average values for five independent calculations in Fig. 10. The non-averaged values of these parameters are shown in Fig. S7. The values of RMSD for the non-hydrogen atoms of the nucleic acids and backbone atoms of the helical-core region of the protein (cdM2-1_75–171_) reached stable levels after 2.5 and 1.0 ns, respectively, whereas for the entire cdM2-1 observed significant differences due to the flexible terminal regions contributions (Fig. 10A). The number of contacts between atoms of the cdM2-1_75–171_ and RNA molecules for distances < 0.6 nm was almost stable around 5200 (Fig. 10B), indicating that the protein and nucleic acids remained interacting throughout the simulation time. Fig. 10C shows that the number of hydrogen bonds formed between the cdM2-1_75–171_ and nucleic acids presented very stable along all the MD simulations with an average value around nine, being Lys150 and Arg151 responsible for at least one hydrogen bond each.

**Figure 10.**
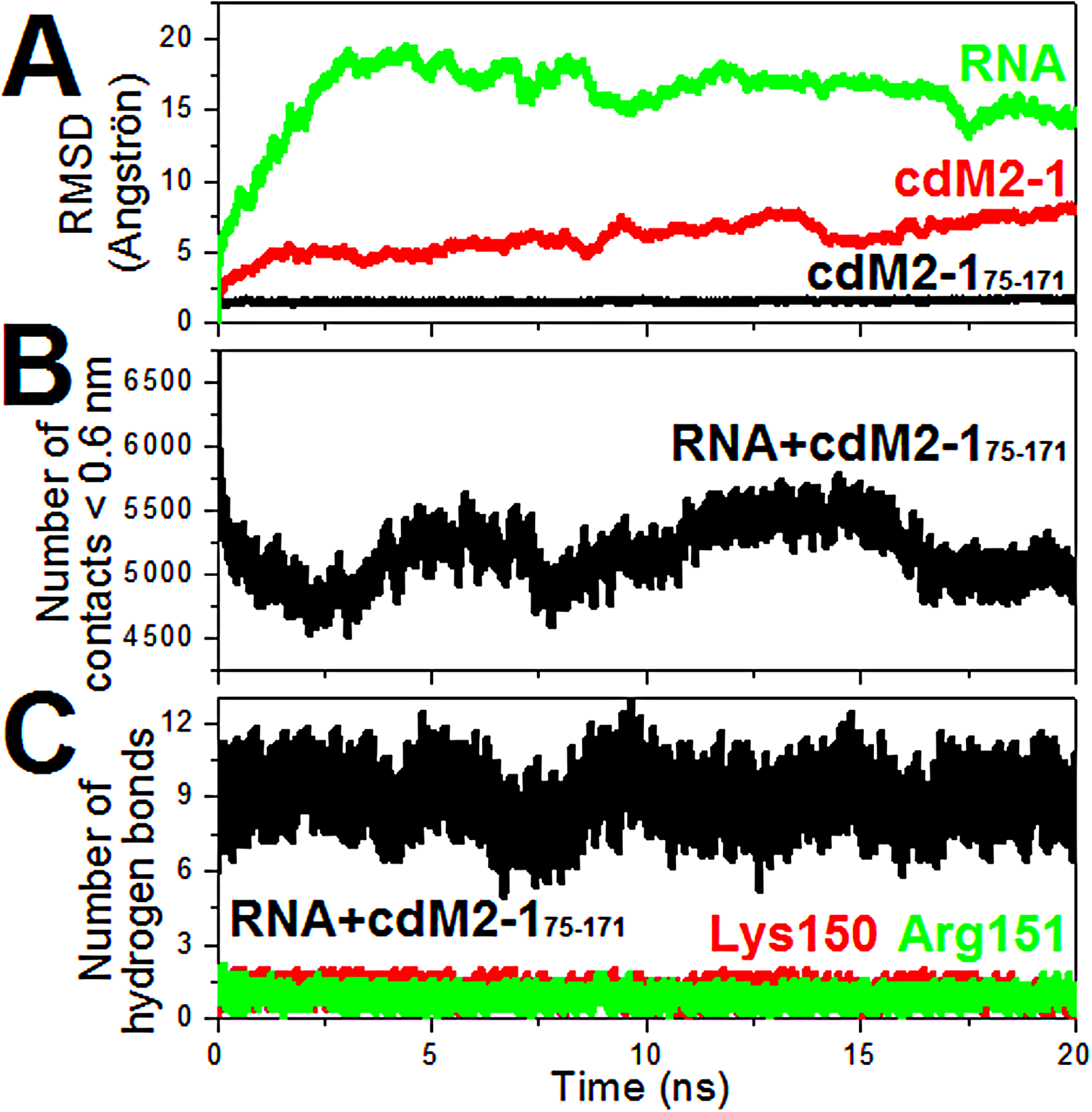
Analysis of the molecular dynamics of the structural models of the cdM2-1/RNA complex. (A) RMSD values for the non-hydrogen atoms of RNAs (green line), for the backbone atoms of the helical-core region of the protein (cdM2-1_75–171_, black line), and for the backbone atoms of the entire cdM2-1 (red line). (B) Number of contacts between atoms of cdM2-1_75–171_ and RNAs for distances < 0.6 nm. (C) Number of hydrogen bonds formed between the RNA molecules and the cdM2-1_75–171_ (black line), the residue Lys150 (red line), and residue Arg151 (green line).

## DISCUSSION

The hRSV M2-1 protein is a transcription antitermination factor that is important for the efficient synthesis of full-length mRNAs as well as for the synthesis of polycistronic read-through mRNAs (11). Molina and collaborators (2018) showed that 20mer RNAs are unfolded upon the formation of its complex with M2-1 tetramer and this finding corroborates with the one of hypotheses proposed by Blondot and colleagues (2012) for the M2-1 activity mechanism either preventing mRNA to re-hybridize to the template or from forming secondary structures. Here it was presented the first evidence that cdM2-1 alone has an unfolding activity for long RNAs and therefore has an active role in the function of the full-length protein as a processivity and antitermination factor of the RdRp complex. The intensity of the imino proton resonances is a direct measure of the base-pairing that stabilize RNA secondary structure. The reduction in these resonance intensities of the protein-bound nucleic acid (Fig. 7) might be explained by the destabilizing activity of the α1–α2–α5–α6 helix bundle of cdM2-1, promoting an unfolding of the RNA secondary structures. This structural finding is different from that reported by Molina and collaborators (2018), in which they observed no significant structural change from the 20mer RNA band at 270 nm in presence of cdM2-1 via CD measurements (10). However, it is worth mentioning that NMR and CD techniques have obviously distinct resolution powers what could explain the observed differences. The evidence that the cdM2-1 alone induced structural changes in long nucleic acid molecules (secondary structure unfolding) may be a consequence of the molecular recognition of RNA base-pairing by this domain. Such logical thinking corroborates with the outcome reported by Blondot and collaborators (2012) in which cdM2-1 presented a higher affinity by double-stranded RNA (12mer) than by its corresponding single-stranded ones, highlighting the key role of base-pairing of the nucleic acid for the cdM2-1/RNA interaction (7).

The fluorescence quenching experiments showed that the binding of the hRSV cdM2-1 to RNA has nano to micromolar dissociation constants (Table 1). Similar values of *K_d_* were reported by Blondot et al. (2012) and Molina et al. (2018) for the binding of the cdM2-1 to nucleic acids with a length between 10 and 20 nucleotides with sequence specificity, being that the first group determined values in 10–100 micromolar range using CSP by NMR spectroscopy and the second, in the hundred-nanomolar range using EMSA (Electrophoretic Mobility Shift Assay) and fluorescence anisotropy (7, 10). Studies reported that the interaction of the full-length M2-1 with nucleic acids present dissociation constants of tenths of nanomolar that are at least 10fold lower than those recorded for the cdM2-1: 30 nM for long RNA without sequence specificity (22), ~19 nM for poly-A 13mer RNA (8), and ~15 nM for 20mer RNA with sequence specificity (10). This indicates that although the core domain is responsible for the first molecular contact with the nucleic acid molecules, there is another region on full-length M2-1 that provides an increase in the affinity with RNA. Results in the literature suggest that these regions may be the zinc finger domains, which provide a type of binding specificity for adenine bases at specific positions in the nucleotide sequence (9), and/or the neighboring core domains in the tetrameric arrangement, promoting a positive cooperative binding to two RNA molecules of 13mer or longer per tetramer of the M2-1 (10).

The effects of temperature, ionic strength, and pH revealed important features of the cdM2-1/RNA interaction. The binding affinity of the cdM2-1 towards the nucleic acid decreased as the temperature increased, indicating that the interaction is enthalpy driven (Δ*H* <0) and entropically unfavorable (Δ*S* <0) (Table 1). The analysis of these thermodynamic parameters pointed out that the stabilization of the complex involves short-range interactions, such as van der Waals forces and hydrogen bonds (13), while the evaluation of the effect of NaCl concentrations and pH changes suggested that the formation of the complex is dependent on the electrostatic interactions for driving the bimolecular encounter between cdM2-1 and RNA. These results are important to understand the M2-1/RNA interaction process since they indicate that this binding process may be tuned by physicochemical conditions of the molecular environment where these biomolecules are, such as might take place in cytoplasmic inclusion bodies in which all the components of the viral RNA polymerase are concentrated, including M2-1, and where the hRSV RNA synthesis occurs (23).

DSC data showed that the melting temperature of the hRSV cdM2-1 (68.7 °C) is at least 13 °C higher than those reported for the full-length M2-1 (~55 °C) (15). Similar results were reported for other proteins in the literature, such as the C-terminal domain (CH2) of the human utrophin calponin-homology domain (24) and the N-terminal domain (C2A) of the cytosolic C2 domains in tandem of synaptotagmin I (25). These findings are especially interesting because isolated domains of multi-domain proteins are usually less stable than the corresponding full-length polypeptide chains. Such a fact can be explained by the lack of favorable interactions with tethered domains in the multi-domain context (26). It is noteworthy that Molina and coauthors observed that the GdmCl unfolding transitions of the core domain is superimposable on that of the M2-1 monomer, being suggested that contacts of the core domains with other core domains or other regions (i.e. the zinc finger region) in the tetramer must be regulatory, with no effect on the conformation or stability of the core domain (27). However, the data presented herein indicate that the thermal stability of the hRSV cdM2-1 is affected due to contacts in the full-length protein which may promote a type of negative inter-domain coupling interaction. The DSC data also showed that values of melting temperature (*T_m_*) of the cdM2-1 had a slight increment due to the interaction with RNA, thus suggesting a slight increase in thermal stability of the protein and also a preferential binding of RNA to the native state of the core domain (28).

The analysis of chemical shift perturbation (CSP) showed that the α1–α2–α5–α6 helix bundle was the main protein region involved in the cdM2-1/RNA interaction interface. The residues taking place on this RNA-binding protein region were similar to those determined by Blondot and collaborators (2012) (7). The α1–α2–α5–α6 helix bundle delimits a positive electrostatic potential surface on cdM2-1 (Fig. 4B) composed of positively charged residues (Lys92, Lys150, Arg151, Lys158, and Lys159) which are responsible for the formation of the complex, especially Lys92, Lys150 and Arg151 that showed large chemical shift changes (Fig. 3B) and also are residues conserved among Respiratory Syncytial Virus and Metapneumovirus M2-1 proteins (Fig. S8). This favorable electrostatic contribution to the interaction with a negative charge of the phosphate groups from nucleic acid corroborates with the analysis of NaCl concentration and pH change effects on the formation of the cdM2-1/RNA complex determined by fluorescence quenching measurements (Fig. 1C and 1D).

The thermal susceptibility measurements of the amide hydrogen (^1^HN) chemical shifts revealed details of structural features of the hRSV cdM2-1 in its free and RNA-bound state. The random coil structure of the N-terminal of cdM2-1 reported by NMR structure (7) may have some level of order since the amide groups of the residues Glu59, Ala63, Asp67, Thr69, Glu70, Glu71, and Ala73 presented a significant propensity to form intramolecular hydrogen bonds (*dδ_HN_/dT* > −5.0 ppb·K^−1^, Fig. 6A). This fact may be explained by propensity of these residues take place in α-helix secondary structure in the full-length hRSV M2-1 protein (Fig. S9). Ile146 in α5-helix showed an unusual *dδ_HN_/dT* value (< −5.0 ppb·K^−1^, Fig. 6A) for a residue in secondary structure, which stands out as a possible point of break in the hydrogen bond network of this helix (green sticks in Fig. 6C). The RNA-bound cdM2-1 revealed changes of *dδ_HN_/dT* values for residues in α1–α2–α5–α6 helix bundle, mapping an interaction interface as that determined by CSP measurements (Fig. 4A), and for residues in α3–α4 hairpin possibly because of a conformational rearrangements remote from the binding region. Two clusters of residues with significant *dδ_HN_/dT* changes (Δ*dδ_HN_/dT* + SD far away from zero), the first in the N-terminal of α2-helix with hydrogen bonds becoming less expandable and the second in N-terminal of α6-helix turning into more expandable, suggest that the main RNA anchor region suffered local structural changes. The side chain of Gln93 presented a decrease in |*dδ_HN_/dT*| (Δ*dδ_HN_/dT* <0) due to the interaction with RNA, indicating that its hydrogen bond became less expandable and possibly stablished it with the nucleic acid since it is not involved in intramolecular hydrogen bonds in NMR structures. On the other hand, the side chain of Asn141 forms intramolecular hydrogen bond with Ser137 in α4–α5 loop of the NMR structures and it became more expandable under binding to RNA (|*dδ_HN_/dT*| increased, Δ*dδ_HN_/dT* >0), possibly because of a conformational rearrangement of the α3–α4 hairpin.

The sPRE experiments revealed that the N and C-terminus of the cdM2-1 and notably the N-terminal residues of the α1-helix are protein regions significantly exposed to the solvent, corroborating with the solution structure of cdM2-1 determined by NMR spectroscopy (7). Unlike the other helices, the residues Ser122, Arg126, Thr130, Ile133, and Ser137 in α4-helix exhibited a nearly solvent-exposed *i* + 4 pattern with comparatively high sPRE effect (Fig. 5), which suggests that the face composed of these residues forms a solvent accessible cleft between α1–α2–α5–α6 helix bundle and α3–α4 hairpin (Fig. S3). For RNA-bound cdM2-1, it was observed a significant reduction of the sPRE effect throughout all non-terminal residues (Fig. 5), indicating that the protein with exception of its N and C-terminus is permanently protected from solvent exposure when it is complexed with nucleic acid. This result suggests that an RNA region anchors mainly in α1–α2 and α5–α6 loops while other non-interacting portions of its length protrude over the α1–α2–α5–α6 helix bundle, wrapping the helical region of cdM2-1.

The ^15^N backbone dynamics data revealed that the N and C-terminal residues of the cdM2-1 have fast motions on pico to nanosecond timescale even after the formation of the complex with RNA, corroborating with sPRE experiments which showed that these residues are solvent exposed. As with CSP and thermal susceptibility measurements, the backbone dynamics data showed that the terminal residues did not take part in the interaction with the nucleic acid. Excluding N and C-terminus, the decrease in R_1_ and increase in R_2_ indicates that the overall tumbling motion of the cdM2-1 decreased upon binding to RNA when compared with free protein in solution and thus confirms the formation of the complex. As the average R_2_/R_1_ ratio is proportional to rotational correlation time (*τ_C_*), R_2_/R_1_ values suggested that apparent *τ_C_* value of the protein increased at least two-fold after the binding to the nucleic acid. The values of R_2_/R_1_ ratio also revealed that residues in α6-helix underwent dynamics on micro to millisecond timescale due to the interaction with RNA, while those residues in α1–α2 loop remained in conformational exchange processes. It is worth pointing out that helices α1–α2 loop and α6-helix cdM2-1 are important for the binding to nucleic acids as it was determined herein by CSP analysis and thermal susceptibility measurements and also reported by Blondot et al. (2012) (7). The extended model-free formalism of Lipari-Szabo of the ^15^N backbone dynamics data of cdM2-1 confirmed that the core domain of hRSV M2-1 is monomeric in solution since the calculated value of *τ_C_* (8.53 ± 0.07 ns) is consistent with the analysis of average R_2_/R_1_ rate and with the tumbling of proteins with a molecular mass of ~14 kDa. The value of below-average order parameter (*S*^2^) along with *dδ_HN_/dT* < −5.0 ppb·K^−1^ for Ile146 in α5-helix reinforced the assumption that this residue is as a point of break of the cdM2-1 secondary structure.

Although raw nuclear spin relaxation data reveal information resulting from the Brownian and internal motion of a protein, interpreting this data to quantify the amplitude and timescales of the dynamics is nontrivial. However, dynamic information within the heteronuclear relaxation rates can be used to evaluate the spectral density functions corresponding to ^1^H-^15^N bonds within a protein to obtain motional features. The mean *J*(0) values of the α2–α3 and α3–α4 loops were slightly lower than the helical-core average for the free protein and, after the binding to nucleic acid, these values significantly decreased with respect to the helical-core average for the RNA-bound cdM2-1 (Table S1). The opposite happened for the mean *J*(0.87*ω_H_*) values of the α2–α3 and α3–α4 loops, these values were slightly higher than the overall average before the formation of the cdM2-1/RNA complex and after that, they significantly increased concerning the helical-core average (Table S1). This inversely correlated behavior between *J*(0) and *J*(0.87*ω_H_*) has been reported by other authors in the literature (29–31) and, in this case indicates that the α2–α3 and α3–α4 loops of the protein became more flexible after the binding to the nucleic acid, since these underwent fast internal motions on the picosecond timescale. This fact can be exemplified for the residue Asn107 in 2– 3 loop and Glu119 in α3–α4 loop (Fig. 8A and 8C). The mean values of *J*(*ω_N_*) spectral density of the α2–α3 and α3–α4 loops were significantly lower than the helical-core average for the RNA-free cdM2-1 and, after the interaction with the nucleic acid, these values approached the overall average (Table S1). The residues Asp110, Lys113 and Lys114 in α3-helix showed a similar behavior to these loops (Fig. 8B). The direct and inverse correlations observed from the plot of *J*(*ω_N_*) versus *J*(0) for the protein in absence and presence of RNA (Fig. S5B and S5D), respectively, suggest that fast internal motions on the picosecond timescale took place in α2–α3 and α3–α4 loops and in α3-helix before complexation and after that, these fast dynamics increased. This result corroborates with the analysis for the spectral density functions *J*(0) and *J*(0.87*ω_H_*).

The molecular docking calculations revealed that the structural models of the cdM2-1/RNA complex predicted by 3dRPC webserver (20) are in agreement with the analysis of CSP and sPRE from NMR data (Fig. 3, 4, and 5), which suggest that despite the nucleic acids specifically bind to the distal portions of the α1, α2, α5, and α6 helices, the RNA molecules may surround the entire helical-core region of the protein (Fig. 9A, 9C, and 9D). The theoretical structural models of the complexes also corroborate with the evidence of the structural changes (unfolding) of RNA induced by cdM2-1 verified by NMR (Fig. 7), since the models indicate that the core domain of hRSV M2-1 preferably binds to secondary structure regions of the nucleic acid where there is base-pairing formation (Fig. 9B). An analysis of non-covalent interactions that stabilize the cdM2-1/RNA binding in the structural models presented the occurrence of hydrogen bonds, salt bridges, and π-cation interactions, what is in line with the interpretation of the thermodynamic parameters from the fluorescence data (Table 1). The molecular dynamics calculations proved the stability of the structural models of the cdM2-1/RNA complex over 20 ns simulations (five independent calculations), revealing a constancy of molecular contacts (~5200), stable values of RMSD after at least 2.5 ns, and an average of nine hydrogen bonds (Fig. 10).

In conclusion, the present work points out the importance of the electrostatic interactions in the formation of the cdM2-1/RNA complex, as well as hydrogen bonds and van der Waals forces in the stabilization of the bimolecular complex, highlighting Lys150 and Arg151 as key residues for the binding which also were identified from mutagenesis assays by Blondot et al. (2012) as crucial for *in vitro* RNA binding to cdM2-1 and for transcription enhancement *in vivo* by full-length M2-1 (7). The α1–α2–α5–α6 helix bundle of the core domain of hRSV M2-1 protein characterizes as the main binding region to the nucleic acid, undergoing local conformational changes upon RNA-binding and also promoting an unfolding of partially structured regions of the long RNAs triggered by the molecular recognition of the base-pairing. It is worth mentioning that this finding that the cdM2-1 alone has an unfolding activity for long RNAs can be essential in the understanding of this activity of the full-length protein, where cooperativity takes place in the M2-1/RNA binding. Although the α3–α4 hairpin is not directly involved in the interaction with the nucleic acid, it shows punctual conformational rearrangements remote from the binding region, as well as an increase in the internal motions on picosecond timescales of the α2–α3 and α3–α4 loops of the cdM2-1. The core domain as a whole is surrounded by the long nucleic acid molecules and the most likely hypothesis is that the α1–α2–α5–α6 helix bundle preferentially binds to the RNA base-pairing while its other regions protrude over the helical-core region of the protein. Therefore, it is revealed that the cdM2-1/RNA complex originates from a fine-tuning binding which likely contributes to the interaction aspects required to processivity and antitermination activity of the M2-1.

## MATERIAL AND METHODS

### Sample preparation

The core domain (residues 58–177) of hRSV M2-1 (cdM2-1) was expressed in *E. coli* BL21 (DE3) RIL cells with a pD441-NHT vector (ATUM, USA) in M9 minimal medium containing ^15^NH_4_Cl as the sole nitrogen source for production of isotopically labeled protein, as described previously (32). The construct of vector includes an N-terminal hexahistidine affinity tag (His6-tag) and a TEV cleavage site. After the expression, the cell suspension was centrifuged and the pellet was resuspended in buffer A (50 mM Tris-HCl pH 8.0, 500 mM NaCl, 1.0 mM β-mercaptoethanol, and 5% (v/v) glycerol). The cells were lysed by sonication and next the crude extract was centrifuged. The supernatant was loaded on a Ni-NTA column for affinity chromatography, pre-equilibrated with buffer A. The column was washed extensively with buffer A containing 10, 20, and 40 mM imidazole, and the protein was eluted with a 60 to 500 mM imidazole gradient. The eluted protein fractions were digested for removing of the His6-tag using TEV protease at 20 °C for 14 hours. This step was performed via dialysis process using a solution containing 20 mM Tris-HCl (pH 8.0), 1.0 mM DTT, and 0.5 mM EDTA. Next, the protein without His6-tag was injected into a Superdex 75 10/300 GL (GE Healthcare) size exclusion column with the buffer B (50 mM NaH_2_PO_4_/Na_2_HPO_4_ pH 6.5, 150 mM NaCl, and 1.0 mM DTT) used for fluorescence quenching, CD, DSC, and NMR experiments. In the fluorescence quenching and CD experiments, the protein sample was dialyzed against a citrate/phosphate and mono/dibasic phosphate buffer to reach the pH 5 and 8, respectively. For NMR measurements, it was added 0.1% (w/v) NaN_3_ and 7% (v/v) D_2_O into phosphate buffer solution (buffer B). The RNA from Torula Yeast type VI (~80 bases) was purchased from Sigma-Aldrich. The stock solution of RNA was prepared at the same buffer solution as for protein, taking into account the pH control because of the solubilization of the nucleic acid. The concentrations of cdM2-1 and RNA were determined by absorbance at 280 and 260 nm using extinction coefficients of 5,960 M^−1^·cm^−1^ (Expasy-ProtParam) (14) and 8,050 M^−1^·cm^−1^·base^−1^ (ThermoFisher Scientific, DNA and RNA Molecular Weights and Conversions), respectively.

### Fluorescence quenching measurements

The fluorescence quenching experiments were performed using Fluorescent Spectrometer Lumina (Thermo Fisher Scientific, USA) equipped with a Peltier system for temperature control and a 10 mm optical path quartz cuvette. The excitation wavelength at 288 nm was set to promote the fluorescence emission of the protein. The emission spectra were reported in a 300– 450 nm range with increment of 1.0 nm, which was corrected for background fluorescence (from buffer) and for inner filter effects (33). Both excitation and emission bandwidths were set at 5 nm. Each point in the emission spectrum is the average of 10 accumulations. The titrations were performed adding small aliquots of the RNA stock solution (715 μM) to 2 mL of cdM2-1 at constant concentration (5.5 μM). The titration experiments were reported at different temperatures (15, 25, and 35 °C) for determining the binding thermodynamic profile of the cdM2-1/RNA complex, as well as at different conditions of salt concentration (0, 150, and 350 mM of NaCl) and pH (5, 6.5, and 8). Measurements were performed in duplicate.

The fluorescence quenching data of the cdM2-1/RNA interaction were analyzed using the following equation (34):

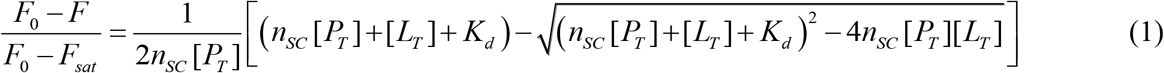

where *F*_0_ is the fluorescence intensity in absence of the ligand (RNA), *F* the fluorescence in presence of RNA, *F_sat_* the intensity of the bound protein saturated with RNA, *K_d_* the dissociation constant, *n_SC_* the stoichiometry coefficient of the cdM2-1/RNA complex corresponding to the number of protein molecules bound to one nucleic acid molecule, [*P_T_*] the total concentration of the protein, and [*L_T_*] the total concentration of the ligand. The values *K_d_* and *n_SC_* were determined from the fitting process to the experimental data by nonlinear least-squares optimization using Levenberg-Marquardt interactions.

### Thermodynamic profile analysis

The driving forces responsible for the binding between biomolecules may include electrostatic interactions, hydrogen bonds, van der Waals interactions, and hydrophobic contacts (13). With the purpose of elucidating the interactions involved in the cdM2-1/RNA complex, the thermodynamic parameters were calculated from the van’t Hoff equation:

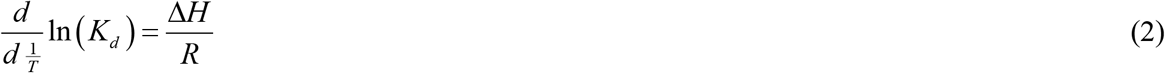

where Δ*H* is the enthalpy change, *R* the universal gas constant, and *K_d_* the dissociation constant at the correspondent temperature (*T* = 15, 25, and 35 °C). The Δ*H* value was obtained from the slope of the van’t Hoff plot, with the respective values of Gibbs free energy changes (Δ*G*) determined from Δ*G* = *RT*ln(*K_d_*) and entropy change (Δ*S*) from Δ*S* = (Δ*H* – Δ*G*)/*T*.

### Circular dichroism measurements

The circular dichroism (CD) experiments were carried out using a Jasco 710 spectropolarimeter (Jasco, USA) equipped with a quartz cell of 0.5 mm optical path length. The far UV-CD spectra of 5.5 μM cdM2-1 were recorded at 25 °C and pHs 5, 6.5, and 8. The spectra were averaged over 20 scans in a 260–200 nm range with a resolution of 0.2 nm at the scan speed of 50 nm·min^−1^ and 1.0 nm spectral bandwidth. The CD signal taken as millidegrees (*θ*) was corrected for the background contribution of the buffer and next expressed in terms of mean residues ellipticity [Θ] in deg·cm^2^·dmol^−1^ using [Θ] = *θ*(*m*deg) /10[*P*]*ln_R_*), where [*P*] is the molar concentration of the protein, *l* is the optical path length (cm), and *n* is the number of amino acid residues.

### Differential scanning calorimetry

Differential scanning calorimetry (DSC) experiments were performed using an N-DSC III (TA Instruments, USA) in the temperature range 10–90 °C at a heating and cooling scan rate of 1.0 °C/min. Both calorimetry cells were loaded with the buffer solution, equilibrated at 10 °C for 10 min and scanned repeatedly as described above until the baseline was reproducible. Next, the sample cell was loaded with 100 μM of cdM2-1 plus 0, 50 and 100 μM of RNA, and scanned. The thermograms of RNA in buffer solution were recorded as a control for the baseline. The baseline corrections were obtained by subtracting the buffer (or RNA) scan from the corresponding protein scan. Measurements were performed in duplicate. The calorimetry enthalpy change (Δ*H_cal_*) of the unfolding process of cdM2-1 was calculated from the area under the thermogram curve. The van’t Hoff enthalpy change (Δ*H_vH_*) of the thermal denaturation process was obtained from Eq. (2), replacing dissociation constant by unfolding constant (*K_U_*). The values of *K_U_* were calculated by *f_U_*/(1 – *f_U_*), where *f_U_* is the fraction of unfolding protein determined from an integral process of the denaturation thermogram.

### Chemical shift perturbation analysis

The chemical shift perturbation (CSP) method was used to map the amino acid residues of cdM2-1 involved in the binding to RNA. The two-dimensional (2D) ^1^H–^15^N HSQC experiments were carried out in increasing amount of RNA from 0 to 115 μM which were added to the [U– ^15^N] cdM2-1 solution at constant concentration of 350 μM. The ^15^N HSQC spectra were recorded at 25 °C on NMR Bruker Avance III spectrometer of 14.1 T operating at a ^1^H frequency of 600 MHz (Bruker BioSpin GmbH, Germany) equipped with cryogenically cooled Z-gradient probe. The data matrix of the spectra consisted of 1024* × 128* data points (were n* refers to complex points) with acquisition times 106.5 ms (*t_HN_*) and 87.6 ms (*t_N_*). A total of 16 scans per complex *t_N_* increment were collected with a recycle delay of 1.25. The resonance assignment of the core domain of hRSV M2-1 was obtained from the repository BMRB (www.bmrb.wisc.edu) from the access code 17451 (32).

The ^1^H-^15^N HSQC spectra were processed using NMRPipe (35) and analyzed using CcpNMR Analysis (36). The chemical shift perturbation of the amino acid residues of the protein recorded in each titration step of RNA against [U–^15^N] cdM2-1 solution was normalized using 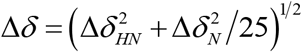, (37), where Δ*δ_HN_* and Δ*δ_N_* denote the chemical shift difference of ^1^H and ^15^N, respectively, recorded in absence and presence of RNA. The mean Δ*δ* of the amino acid residues as well as its standard deviation of CSP were used as cutoff value to identify the residues of the protein involved in the cdM2-1/RNA interaction.

### Amide hydrogen chemical shift temperature coefficient

The amide hydrogen (^1^H_N_) chemical shift temperature coefficient of cdM2-1 was determined by recording a series of 2D ^1^H–^15^H HSQC spectra at 15, 20, 25, 30, and 35 °C in absence and presence of 115 μM RNA, using a NMR Bruker Avance III spectrometer of 14.1 T operating at a ^1^H frequency of 600 MHz (Bruker BioSpin GmbH, Germany). All spectra were referenced to the water signal for each temperature, next processed using NMRPipe (35) and analyzed using CcpNMR Analysis (36). The chemical shift values (*δ_HN_*) of all residues at different temperature were plotted as a function of temperature. The resulting slope (*dδ_HN_/dT*) of every curve was plotted for each residue. For evaluation of the *dδ_HN_/dT* values, it was used a straightforward interpretation compilation published by Morando et al. (2019) (38). In principle, ^1^H_N_ chemical shift temperature coefficient reports on the thermal susceptibility of the H_N_-C′ amide hydrogen bonds, since *dδ_HN_/dT* is correlated to the length of the hydrogen bond (*r_HNC′_*) and hydrogen bond J coupling 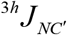 (16). On the one hand, residues with *dδ_HN_/dT* < −5.0 ppb·K^−1^ form more expandable hydrogen bonds, which may be interpreted as weaker since they present smaller hydrogen bond J coupling 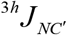. For this reason, an amide group in the protein structure with *dδ_HN_/dT* < −5.0 ppb·K^−1^ may be considered a weakness point of a secondary structure or, when this amide in a loop or intrinsically disordered region (IDR), more exposed to a hydrogen bond with water. On the other hand, residues with *dδ_HN_/dT* > −5.0 ppb·K^−1^ make less expandable hydrogen bonds, which may be interpreted as stronger since they present larger hydrogen bond J coupling 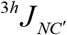. In this sense, an amide group with *dδ_HN_/dT* > −5.0 ppb·K^−1^ is involved in secondary structures of hydrogen bonds or, when this amide in a loop or IDR, may tend to form intramolecular hydrogen bonds and consequently to increase order (16, 38).

### Paramagnetic relaxation enhancement experiments

To monitor the solvent accessibility of residues of cdM2-1 (350 μM) in absence and presence of 115 μM RNA, it was used the gadolinium-based paramagnetic relaxation agent Gd-diethylenetriamine pentaacetic acid-bismathylamide (Gd-DTPA) (GE Life Science, United Kingdom). The ^1^H_N_/^15^N amide cross-peak intensities of the protein were determined by recording a series of 2D ^1^H–^15^H HSQC spectra at different concentration of Gd-DTPA: 0, 1.0, 2.0, 3.0, and 4.0 mM at 25 °C using a NMR Bruker Avance III spectrometer of 14.1 T operating at a ^1^H frequency of 600 MHz (Bruker BioSpin GmbH, Germany). All spectra were processed using NMRPipe (35) and analyzed using CcpNMR Analysis (36). The intensities (*I_NH_*) of each ^1^H_N_/^15^N cross-peak of the residues were plotted as a function of Gd-DTPA concentration and the slope of the adjusted straight line (*I_NH_* · [*Gd*]^−1^) were plotted for each residue. To calculate an accurate slope *I_NH_* · [*Gd*]^−1^, the points where the intensities are close to zero (lack of linearity) were not taken into account.

### Imino proton resonances of RNA

The unidimensional ^1^H excitation sculpting spectra were collected at 25 °C on NMR Bruker Avance III spectrometer of 14.1 T operating at a ^1^H frequency of 600 MHz (Bruker BioSpin GmbH, Germany) using 180° water-selective pulse of 2 ms for the solvent suppression. The free induction decays (FIDs) were recorded with 32,768 data points using a spectral width of 21,000 Hz (35 ppm), a relaxation delay of 1.25 s, and an acquisition time of 0.78 s. The experiment was performed for the free protein at 350 μM and the free nucleic acid at 115 μM and after that, for the cdM2-1/RNA complex maintaining the same concentrations as before. The spectral region from 10 to 18 ppm was used for analyzing structural changes of RNA induced by cdM2-1, probing the imino protons of the nucleic acid involved in the base pair formation (17).

### NMR relaxation experiments

The backbone dynamics of the [U–^15^N] cdM2-1 at 350 μM and the core domain complexed with 115 μM of RNA were investigated from ^15^N nuclear spin relaxation experiments (39) at 25 °C by using NMR Bruker Avence III of 14.1 T operating at a ^1^H frequency of 600 MHz (Bruker BioSpin GmbH, Germany). The *R*_1_ experiments were collected using 14 delay times of inversion recovery of 54, 104, 204, 304, 404 (twice), 604, 804, 904 (twice), 1204, 1504 (twice), and 1804 ms. The *R*_1*ρ*_ experiments were performed according to Korzhnev et al. (2002) (40) with 14 spin-lock periods of 10, 20, 30 (twice), 40, 50 (twice), 60, 70, 80, 90 (twice), 100, and 110 ms, and ^15^N spin-lock field strengths of 2.0 kHz. A recycle delay of 3.0 s was used for the *R*_1_ and *R*_1*ρ*_ experiments. The values of *R*_1_ and *R*_1*ρ*_ were determined from non-linear least-square fittings of the intensities using two-parameter mono-exponential equations. *R*_2_ values were determined from the measured *R*_1_ and *R*_1*ρ*_ rates (41). The reported errors for the rates were calculated from the estimated uncertainties for the relaxation delay duplicates. {^1^H}–^15^N steady-state heteronuclear nuclear Overhauser effects (NOE) were measured from pairs of interleaved spectra recorded with (NOE) and without (control) H^N^ proton saturation during a recycle delay of 12 s. The {^1^H}–^15^N hetNOE values were calculated from resonance intensity ratios obtained from the NOE and control spectra, with uncertainties estimated from the background noise of the spectra. The data were processed using NMRPipe (35) and analyzed using CcpNMR Analysis (36).

### Reduced spectral density mapping approach

The reduced spectral density mapping (rSDM) is a simplified approach for analysis of nuclear spin relaxation data (R_1_, R_2_ and hetNOE) developed by Farrow et al. (1995) (42), Ishima & Nagayama (1995) (43), and Lefèrve et al. (1996) (44). The rSDM maps the spectral density for determining the accurate values of the spectral density function at three frequencies: *J*(0) at the zero frequency, *J*(*ω_N_*) at the nitrogen frequency, and 〈*J*(*ω_H_*)〉 at an effective proton frequency. The approach exploits the assumption that at higher frequencies, the spectral density terms contributing to the relaxation processes are equal in magnitude: *J*(*ω_H_*) ≈ *J*(*ω_H_* ± *ω_N_*); and is replaced by a single equivalent term 〈*J*(*ω_H_*)〉, which corresponds to *J*(0, 87*ω_H_*). This assumption is based on the premise that at higher frequencies in comparison with *J*(0) and *J*(*ω_N_*), spectral densities *J*(*ω_H_*) e *J*(*ω_H_* ± *ω_N_*) show minimal variation. The spectral density functions at three frequencies, as derived through reduced spectral density mapping, are given by the following equations:

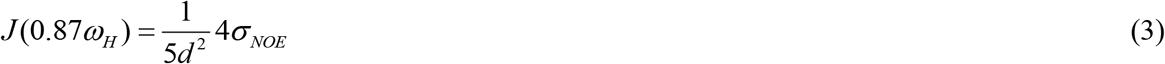

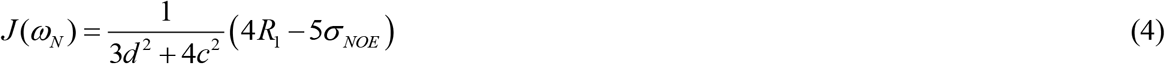

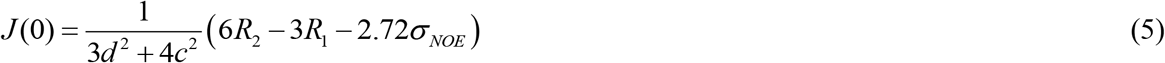

where the cross-relaxation rate is defined by 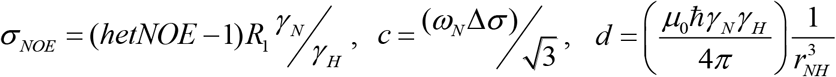, *μ*_0_ is the permeability of the vacuum, *ℏ* is reduced Planck constant, *γ_N_* and *γ_H_* are the gyromagnetic ratios of ^15^N and ^1^H, respectively, *r_NH_* is the bond length, *ω_N_* is the Larmor frequency of the ^15^N nucleus, and Δ*σ* is the ^15^N chemical shift anisotropy in ppm.

### Model free analysis

The relaxation parameters were fitted according to the extended model-free formalism of Lipari-Szabo for obtaining the intramolecular dynamics (45). TENSOR2 program (18) was employed to define a motional model for hRSV cdM2-1 using an anisotropic tensor with a fully asymmetry diffusion model, (*D_X_* = 1.81 ± 0.03, *D_Y_* = 1.95 ± 0.02, *D_Z_* = 2.11 ± 0.03) × 10^7^ s^−1^. The calculations of the rotational correlation time (*τ_C_*) were performed from the relaxation data for the amino acid residues with values of R_2_/R_1_ ratios within one standard deviation relative to the calculated average and with values of {^1^H}–^15^N NOE heteronuclear higher than 0.65. The anisotropic diffusion tensor also was used for calculating the internal molecular dynamics parameters, such as order parameter (*S*^2^) and conformational exchange rate (*R_ex_*), which reflects movements of the NH bond in pico to nanosecond and micro to millisecond timescales, respectively.

### Charge, protonation state, and electrostatic potential calculation

The electrostatic potential calculations were performed by the APBS software (46) using charge values and protonation states obtained from the PDB2PQR server (version 2.1.1) (47) along with the PROPKA program (version 3.0) (48). The physical-chemical parameters used for the calculations were 150 mM NaCl, pH 6.5, and 25 °C. The electrostatic potential surface of the core domain of hRSV M2-1 was displayed using PyMOL (49).

### Molecular modeling and molecular docking

The primary sequences of 10 RNAs were generated from the Random Sequence Generator (RSG) webserver (www.molbiotools.com) setting the length of sequence equal to 40 and AU content at 80%. All RNA sequences generate by RSG server are presented in Table S13. These random RNA sequences were submitted to Direct Coupling Analysis (DCA) server (www.biophy.hust.edu.cn) which is a statistical inference framework used to infer direct co-evolutionary couplings among nucleotide pairs in multiple sequence alignments, which aims at disentangling direct from indirect correlations. The information provided by DCA server was subsequently used in 2dRNA webserver (www.biophy.hust.edu.cn) which is a secondary structure prediction method of RNA based on DCA data. The parameters of 2dRNA server were set as default. Next, the predicted secondary structure information was used in 3dRNA webserver (version 2.0) (20) which is automatic molecular modeling method for building the three-dimensional structure of RNA. The assembly calculations performed by 3dRNA server were followed by an optimization step using all the parameters as default.

The molecular docking calculations for predicting the molecular model of the complex formed between the random RNA sequences (Table S13) and cdM2-1 were performed by using the 3dRPC webserver (version 2.0) (50). The three-dimensional structure of the hRSV M2-1 core domain was downloaded from the Protein Data Bank, access code 2L9J (7), and 3D structures of 10 RNAs were obtained from 3dRNA webserver (version 2.0) (20). A total of 10 complex models were predicted using the 3dRPC-Score function. The amino acid residues of cdM2-1 involved in the RNA binding interface, which was determined from CSP method by NMR experiments, were specified as a constraint set in the advanced setting of the 3dRPC server. Following docking, the lowest energy structural models of 10 cdM2-1/RNA complexes were analyzed by VMD software (51) using the VolMap tool for evaluating of the mass density map of collocated RNA (1.0 Å of resolution; 1.5 Å × radius of atom size; 10 frames combined using average; and isosurfaces with isovalue of 0.5) and by PLIP webserver (21) for characterizing the protein/nucleic acid non-covalent interactions, such as hydrogen bond, π-cation interaction, π-stacking, and salt bridge. Structural conformation of the constructed models was displayed using PyMOL (49) and VMD (51).

### Molecular dynamics simulation

The molecular dynamics (MD) simulations were carried out with the GROMACS program (version 5.0.7) (52). The molecular systems were modeled by using the AMBER99SB-IDLN protein and AMBER94 nucleic acid force field (53), and TIP3P water model (54). The three-dimensional structures of 5 random-selected cdM2-1/RNA complexes calculated from molecular docking (3dRPC server) were used in the MD simulations. These structures were placed in the center of 89–135 Å cubic boxes filled with TIP3P water molecules and Na^+^/Cl^−^ ions ([NaCl] = 150 mM). The protonation states of charged residues were set considering a pH 6.5 from the PROPKA results (48). All simulations were performed in NPT ensemble using periodic boundary conditions and keeping the system at 298 K (Nose-Hoover thermostat, *τ_T_* = 2.0 ps) and 1.0 bar (Parrinello-Rahman barostat, *τ_P_* = 2.0 ps and compressibility = 4.5×10^−5^ bar^−1^). Lennard-Jones and Coulomb potentials were applied using a cutoff distance of 12 Å. The long-range electrostatic interactions were calculated using the particle mesh Ewald (PME) algorithm. The covalent bonds involving hydrogen atoms were constrained to their equilibrium distance. A conjugate gradient minimization algorithm was utilized to relax the superposition of atoms generated in the box construction process. The energy minimizations were performed with steepest descent integrator and conjugate gradient algorithm, using 500 kJ·mol^−1^·nm^−1^ as maximum force criterion. 100 thousand steps of molecular dynamics were performed for each NVT and NPT equilibration, applying force constants of 1000 kJ·mol^−1^·nm^−2^ to all heavy atoms of the cdM2-1/RNA complexes. Lastly, MD simulations of 20 ns were accomplished for data acquisition. Following the simulations, the trajectories were concatenated and analyzed according to the number of hydrogen bonds (cutoff distance = 3.5 Å and maximum angle = 30°), number of contacts (< 0.6 nm), and root-mean-square deviation (RMSD) of non-hydrogen atoms for the nucleic acid and backbone atoms for cdM2-1 (entire protein and helical-core region). These parameters (hydrogen bonds, number of contact, RMSD) from 5 MD simulations were presented as averages.

### Accessible surface area calculations

The accessible surface areas (ASA) of free cdM2-1 and its docked complex with RNAs were calculated using the NACCESS program (55). The structural models of the cdM2-1/RNA complexes were obtained from the molecular docking calculations (3dRPC server). Changes in absolute ASA for residue *i* were calculated using 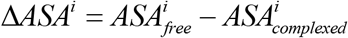, where 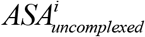 and 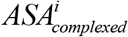 is the absolute accessible surface area for free and RNA-bound cdM2-1 residues, respectively. The values of Δ*ASA* were averaged and next the average value was normalized to a range between 0 and 1.0 units for coloring the cdM2-1 structure.

## ACKNOWLEDGMENTS

The author I.P.C. gratefully acknowledges the financial support by postdoctoral fellowship from FAPERJ (202.279/2018) and the PROPe UNESP. G.C.G., V.B.M. and F.P.S. thank to the FAPESP scholarship (2019/08739-1, 2018/08900-4). The authors thank Prof. Dr. Marcelo de Freitas Lima for the access to the Fluorescent Spectrometer Lumina located in the Bio-organic Environmental Laboratory of the Department of Chemistry and Environmental Sciences, and Prof. Dr. Alexandre Suma de Araújo for the access to the Calix cluster (FAPESP 2010/18169-3) for performing the molecular dynamics simulations. The authors also recognize GridUNESP for the availability of the software package for molecular dynamics simulations. The authors thank for the access to the NMR laboratory of Multiuser Center for Biomolecular Innovation (FAPESP 2009/53989-4) in IBILCE/UNESP. The authors thank NMRbox: National Center for Biomolecular NMR Data Processing and Analysis, a Biomedical Technology Research Resource (BTRR), which is supported by NIH grant P41GM111135 (NIGMS).

